# Medial temporal lobe atrophy patterns in early- versus late-onset amnestic Alzheimer’s disease

**DOI:** 10.1101/2024.05.21.594976

**Authors:** Anika Wuestefeld, Alexa Pichet Binette, Danielle van Westen, Olof Strandberg, Erik Stomrud, Niklas Mattsson-Carlgren, Shorena Janelidze, Ruben Smith, Sebastian Palmqvist, Hannah Baumeister, David Berron, Paul A. Yushkevich, Oskar Hansson, Nicola Spotorno, Laura EM Wisse

## Abstract

**Background:** The medial temporal lobe (MTL) is hypothesized to be relatively spared in early-onset Alzheimer’s disease (EOAD). Yet, detailed examination of MTL subfield volumes and drivers of atrophy in amnestic EOAD is lacking.

**Methods:** BioFINDER-2 participants with memory impairment, abnormal amyloid-β status and tau-PET were included. Forty-one EOAD individuals aged ≤65 years and, as comparison, late-onset AD (LOAD, ≥70 years, n=154) and Aβ-negative cognitively unimpaired controls were included. MTL subregions and biomarkers of (co-)pathologies were measured.

**Results:** AD groups showed smaller MTL subregions compared to controls. Atrophy patterns were similar across AD groups, although LOAD showed thinner entorhinal cortices compared to EOAD. EOAD showed lower WMH compared to LOAD. No differences in MTL tau-PET or transactive response DNA binding protein 43-proxy positivity was found.

**Conclusions:** We found in vivo evidence for MTL atrophy in amnestic EOAD and overall similar levels to LOAD of MTL tau pathology and co-pathologies.

## Background

Early-onset Alzheimer’s disease (EOAD) is commonly defined by a clinical onset before the age of 65 years and is one of the most common types of early-onset neurodegenerative dementia (1). It shares the presence of main neuropathological features, i.e., fibrillar amyloid-β (Aβ) and hyperphosphorylated tau, with late-onset LOAD (age>65), but clinical features and other characteristics tend to differ between EOAD and LOAD (1). For example, there is evidence for less semantic memory impairment and a more aggressive course with more neurofibrillary tangle (NFT) pathology in EOAD compared to LOAD (1,2).

While prior research has investigated clinical, genetic or pathological differences in EOAD vs. LOAD, for example (3–6), many studies define EOAD only by age of onset. Thus, various clinical phenotypes, such as amnestic or non-amnestic EOAD, posterior cortical atrophy (PCA), or primary progressive aphasia (PPA) (7), have been grouped together as EOAD group as they are more common in younger age than for late-onset AD (1). Due to this grouping, observed differences between EOAD vs. LOAD may not be applicable to all clinical phenotypes. For example, the medial temporal lobe (MTL) has previously been found to be relatively spared in EOAD compared to LOAD in several studies (8–10). However, it is unclear if this applies to amnestic EOAD given the common grouping of clinical phenotypes. Moreover, fine-grained changes in MTL subfield atrophy patterns have not been investigated. MTL subfields are heavily involved in memory function (11) but subserve different functions (12,13). Additionally, the cytoarchitectonic and functionally different MTL subfields are differently affected in AD and other neurodegenerative diseases (14–16). The involvement of the MTL in amnestic EOAD is not well characterized, therefore it is of importance to investigate whether the MTL is affected in EOAD and to what extent the atrophy pattern differs from the more common amnestic LOAD (17).

In addition to Aβ and NFT, co-pathologies are also common in AD (18) and can affect the clinical course of the disease as well as atrophy patterns in the brain (18–20). Common AD co-pathologies, such as cerebrovascular disease (CVD) or transactive response DNA binding protein 43 (TDP-43) pathology often occur in the MTL (21,22). Therefore, MTL atrophy patterns in amnestic AD are likely partially influenced by the presence of such co-pathologies. It has been suggested that co-pathologies are common in EOAD, albeit less than in LOAD, and contribute substantially to cognitive impairment in EOAD (5). However, it is unclear if this equally applies to all the phenotypes of EOAD including amnestic EOAD.

In this cross-sectional study we aim to investigate if MTL atrophy occurs in individuals with amnestic early-onset cognitive impairment (EOAD). To this end, we aim to compare MTL subfield differences across amnestic EOAD with the LOAD group and with cognitively normal controls as reference. Secondary aims include (I) investigating similar comparisons for neocortical composite regions in order to establish whether potential differences between EOAD and LOAD groups are specific to the MTL, and (II) assessing if common co-pathologies are present in EOAD vs. LOAD, and in comparison to healthy controls. Lastly, we explore if MTL atrophy is associated with AD pathologies and co-pathologies in the amnestic EOAD group. Exploratory analyses focus on (I) cognitive performance in amnestic EOAD and (II) comparisons with non-amnestic EOAD and LOAD groups.

## Methods

### Participants

We included 534 cognitively impaired from a memory clinic setting and unimpaired participants from population-based studies in the city of Malmö (23) older than 50 years from the Swedish BioFINDER-2 study (NCT03174938) who underwent magnetic resonance imaging (MRI) and tau-PET. The study was approved by the ethical review board in Lund, Sweden, and all study participants provided written informed consent.

Inclusion criteria for the EOAD group were (I) mild cognitive impairment (MCI, MMSE≥24) or AD (MMSE≥20; see details in (23)), (II) 50-65 years of age, and who (III) were Aβ and tau positive accordingly to cerebrospinal fluid (CSF) Aβ42/Aβ40 ratio and tau-PET respectively, and (V) performed 1.5 standard deviations below age- and education-based norms on the Alzheimer’s Disease Assessment Scale-Cognitive subscale (ADAS-cog) delayed word list recall (24). Additionally, patients between 65-70 years of age, who indicated their age of onset was before 65 on the Cognitive Impairment Questionnaire (CIMP-QUEST) and fulfilled all the other criteria were included as EOAD. The LOAD group included only patients with age ≥70 years while the other criteria were shared between EOAD and LOAD. The gap of five years between EOAD and LOAD was chosen to minimize the possibility that EOAD cases were included in the LOAD group. Additionally, in secondary analyses, we included non-amnestic EOAD (naEOAD) and LOAD (naLOAD) participants that had the same group definitions as EOAD and LOAD except that the non-amnestic groups performed within age- and education-based norms on the episodic memory test. We focused only on cases who were Aβ- and tau-positive to ensure that the observed memory or cognitive impairments were at least partly due to AD proteinopathies.

Two control groups were included, one for EOAD and one for LOAD, given the inherent age differences between the patient groups. The control groups were (I) cognitively unimpaired, (II) Aβ negative, (III) performed within age- and education-based norms on the ADAS-cog delayed word list recall, and (IV) were selected with the same age range as respective EOAD or LOAD group.

### Cerebrospinal fluid biomarkers

For a majority of the participants (n=514), CSF levels of Aβ42 and Aβ40 were measured with the Roche Elecsys platform (Roche Diagnostics International Ltd., Basel, Switzerland) as described previously by Hansson et al. (25). For the remaining participants (n=11), Lumipulse G (Fujirebio, n=9) or Meso-Scale Discovery (MSD; n=2) assays, were used to quantify concentration of Aβ42 and Aβ40. All CSF handling followed a standardized protocol (26,27). To determine Aβ-positivity a cut-off for CSF Aβ42/Aβ40 ratio was used with previously described thresholds obtained using Gaussian Mixture Modeling (Elecsys: 0.080; Lumipulse G: 0.072; MSD: <0.077) (28–30).

### Cognitive assesment

Participants’ cognitive functioning was estimated with the Mini-Mental State Examination (MMSE) (31), the Alzheimer’s Disease Assessment Scale-Cognitive Subscale (ADAS-Cog) delayed word list recall (24), animal fluency (32), Boston Naming Test-15 (BNT-15) (33), Trail-Making Test B (34), Symbol digit modalities test (35), and the visual object and space perception (VOSP) battery subtest cubes (36). The scores were z-transformed using Aβ-cognitively unimpaired individuals under the age of 40 from BioFINDER-2 (n=99; MMSE>=26). These cognitive measures were chosen in order to capture various aspects of human cognition, such as memory, visuospatial functioning, language, and processing speed.

### Imaging protocol

#### MRI

T1-weighted, T2-weighted, and Fluid attenuated inversion recovery (T2-weighted FLAIR) images were acquired on a Siemens MAGENTOM Prisma 3T scanner (Siemens Healthineers, Erlangen, Germany) with a 64-channel head coil. Whole brain T1-weighted images (Magnetization Prepared – Rapid Gradient Echo, MPRAGE) were acquired with the following parameters: in-plane resolution=1×1 mm^2^, slice thickness=1 mm, repetition time (TR)=1900 ms, echo time (TE)=2.54 ms, flip-angle=9. Coronal T2-weighted images were acquired using a turbo spin echo sequence (in-plane resolution=.4x.4 mm^2^, slice thickness=2 mm, TR=8240 ms, TE=52 ms, flip-angle=150°) with hippocampal orientation. Similarly, axial T2-weighted FLAIR images were acquired (TR = 5000 ms, TE = 393 ms, TA = 4:37 min with the same resolution and field of view of the T1-weighted images).

#### Structural MRI processing and analysis

Using the Automated Segmentation of Hippocampal Subfields (ASHS) packages for T1- and T2-weighted MR images (37–40), MTL subregions were automatically segmented. To obtain hippocampal subfield volumes (Subiculum, cornu ammonis (CA) 1, dentate gyrus (DG)) the T2-weighted package was used (40). Anterior and posterior hippocampus (HC), and MTL cortical thickness measures (entorhinal cortex (ERC), Brodmann area (BA) 35 (≈transentorhinal cortex), BA36, and parahippocampal cortex) were extracted using the T1-weighted MRI package. Whole amygdala volumes were extracted using ASHS from a new atlas for T1-weighted MRI updated with an amygdala label created following a newly developed protocol (see supplementary methods, sFig. 1-10, sTable 1-2). Volumes of hippocampal subregions and the amygdala were corrected for ICV using volume-to-ICV fractions.

De Flores and colleagues (41) suggested that the ratio between anterior HC and parahippocampal cortex (measured with T1-ASHS) as a promising marker to assess the presence of TDP-43 pathology in dementia cases with AD neuropathologic change and was previously validated against post-mortem data. They propose a cut-off of 693.44 for this marker, indicating the presence of TDP-43 pathology for individuals with a ratio below this cut-off. Following their approach, a ratio between anterior HC volume and parahippocampal cortical thickness was calculated after regressing out ICV for anterior HC and age for both measures and the above-mentioned cut-off was applied.

After applying FreeSurfer 6 (https://surfer.nmr.mgh.harvard.edu/) to the T1-weighted image to obtain mean cortical thickness estimates, the neocortex was parcellated into five composite regions based on the Desikan-Killiany atlas. Average cortical thickness was extracted from the five composite regions consisting of: the lateral temporal (superior, middle, and inferior temporal, banks of the superior temporal sulcus, transverse temporal, temporal pole), lateral parietal (postcentral, inferior and superior parietal, supramarginal), medial parietal (paracentral, isthmus, posterior cingulate, precuneus), frontal (superior frontal, rostral and caudal middle frontal, pars opercularis, pars triangularis, pars orbitalis, lateral and medial orbitofrontal, precentral, paracentral, frontal pole), and occipital (cuneus, lateral occipital, lingual, pericalcarine) cortices.

As supplementary analyses, the Longitudinal Early-onset Alzheimer’s Disease Study (LEADS) signature mean thickness, comprising primarily temporal and parietal regions, was calculated, see (42), and compared between groups.

All regions of interest were averaged across hemispheres. All regions of interest were z-scored to facilitate comparisons between the measures using Aβ-cognitively unimpaired individuals under the age of 40 from BioFINDER-2 (n=99; MMSE>=26) as reference group.

### [18F]RO948 tau-PET

Tau-PET scans were acquired with a digital GE Discovery MI Scanner (General Electric Medical Systems). Tau-PET was performed 70-90 minutes post-injection of ∼370 MBq of [^18^F]RO948. Details of the PET reconstruction have been published previously (43). The Swedish Medical Products Agency and the local Radiation Safety Committee at Skåne University Hospital, Sweden approved the PET imaging.

#### Tau-PET processing and analysis

Standardized uptake value ratios (SUVR) were calculated using an inferior cerebellar reference region for [^18^F]RO948-PET (tau-PET) (44). Using the geometric transfer matrix method (45), partial volume correction (PVC) was performed. See Leuzy et al. (43) for a detailed description of our processing pipeline.

[18F]RO948-PET positivity was defined using a previously defined cut-off of a SUVR>1.362 (43) based on Gaussian Mixture Modeling in the temporal meta-ROI corresponding to Braak I-IV (46).

Tau-PET uptake was measures in two early regions (I) a composite MTL region from ASHS comprising ERC and BA35 from ASHS and (II) the amygdala from ASHS. The decision to use only ERC and BA35 was based on two aspects: (I) it reduces the potential bias caused by off target binding that typically occur around the hippocampus, (II) ERC and BA35 typically show the earliest accumulation of cortical tau pathology (14). Using clusters previously defined with an event-based modelling (EBM) approach, see (47), tau-PET composite measures were calculated for four EBM-based regions of interest (lateral temporal, parietal, frontal, occipital/motor), that match the neocortical composite regions. Lastly, a composite tau-PET SUVR was calculated for the LEADS signature (42).

#### White matter hyperintensity volume processing and analysis

Using FreeSurfer 7.2 Sequence Adaptive Multimodal SEGmentation (SAMSEG) functionality (48,49), white matter hyperintensities (WMH) were segmented from the T2-weighted FLAIR sequence. Whole brain WMH volumes were calculated per participant, corrected for ICV (using volume-to-ICV fractions) and log-transformed. This measure was used for primary analyses. Due to the distribution of the data (many participants with very low values), WMH volumes were also split into low/high based on median-split and used in sensitivity analyses.

### Statistical analyses

Analyses were performed in R 4.0.2 (50). All p-values were controlled for the false discovery rate (FDR, Benjamini–Hochberg procedure). P-values were considered statistically significant at p< 0.05. Group comparisons did not by default include age as a covariate, since the AD groups are defined by age. Only comparisons between controls and respective AD groups included age as covariate in sensitivity analyses.

Differences in demographic variables were tested using t-tests or chi-square tests. We examined group differences between EOAD and LOAD with respective controls and with each other for demographics and cognitive measures.

For our main aim, we examined group differences between EOAD and LOAD with respective controls and with each other for volume/thickness of the MTL regions of interest (3 comparisons) using one-way ANCOVAs along with post-hoc Tukey’s HSD Test for multiple comparisons, including sex as covariate. We also investigate the interaction between age group (young vs. old) and diagnosis (CU vs. AD) in a linear regression model for each region in order to investigate if morphological metrics (i.e. volume or thickness) are differently affected by aging and disease state.

In addition, we characterized the EOAD and LOAD groups further by examining group differences between EOAD and LOAD with respective controls and with each other. This analysis was conducted, first, for the thickness of neocortical composite regions. We used ANCOVA to investigate group differences and performed linear regression models for each region with the interaction between age and diagnosis. Second, groups were compared for all biomarkers of AD- and co-pathologies. We used ANCOVA for continuous outcomes and logistic regression for categorical variables to assess group differences for the positivity on the aHC/PHC ratio (MRI-based proxy for potential TDP-43 positivity), as well as binarized WMH volume (low vs. high). In all analyses, sex was included as covariate.

As sensitivity analyses, age was included as covariate for comparisons between AD groups and controls. Second, for comparisons of AD- and co-pathologies, we included also CSF Aβ42/Aβ40 ratio as a covariate to investigate if differences between all group comparisons were influenced by Aβ. Additional analyses additionally investigated group differences between EOAD and LOAD for the both LEADS signature thickness and tau-PET SUVR.

#### Secondary analyses

As exploratory analyses, we aim to investigate if different pathologies could explain lower region of interest volume/thickness within the EOAD group. To this end, we performed linear regressions predicting region of interest volume/thickness using biomarkers of AD- and co-pathologies including age and sex as covariates.

We explored group comparisons for cognitive performance (ADAS-cog delayed word recall, animal fluency, trail-making test B, VOSP cube, BNT-15). ANOVAs were used including sex and education as covariates. We also explore if differences in volume/thickness were associated with cognitive performance within the EOAD group, including education level, age, and sex as covariates.

In a final step, we also explored group comparisons for the amnestic and non-amnestic EOAD and LOAD groups.

## Results

### Demographics

The whole sample consisted of 534 older adults (56.9% female, mean age 69.2, mean education 12.8 years, 47.4% were *APOE*-ɛ4 carriers). The demographics of the EOAD (n=41) and LOAD (n=154) groups as well as the two control groups are shown in Table 1. Comparing EOAD vs. LOAD, no differences in sex, education, or *APOE* status were observed. A significant difference between LOAD and respective controls was observed for sex (lower proportion of males in the AD groups), and, as expected, *APOE* status (higher proportion of *APOE-*ɛ*4* carriership in the AD groups). There was no difference in diagnosis between EOAD and LOAD groups. Despite selecting AD patients and controls from the same age range, age was significantly, but marginally, different between AD groups and the respective controls, likely due to non-normal distributions within the AD groups. While the age difference is likely negligible, we did adjust for age in sensitivity analyses when comparing the AD groups to their respective control groups.

**Table 1.** Characteristics of the sample.

|  | YCU | OCU | EOAD | LOAD | Total | p-value<br>YCU-EOAD | p-value<br>OCU-LOAD | p-value<br>EOAD-LOAD |
| --- | --- | --- | --- | --- | --- | --- | --- | --- |
| **N** | 188 | 151 | 41 | 154 | 534 | - | - | - |
| **Diagnosis** |  |  |  |  |  | - | - | .713 |
| CU | 188 (100) | 151 (100) | 0 (0) | 0 (0) | 339 (63.5) |  |  |  |
| MCI | 0 (0) | 0 (0) | 16 (39.0) | 65 (42.2) | 81 (15.2) |  |  |  |
| AD | 0 (0) | 0 (0) | 25 (61.0) | 89 (57.8) | 114 (21.3) |  |  |  |
| **Sex (female)** | 103 (54.8) | 99 (65.5) | 20 (48.8) | 82 (53.2) | 304 (56.9) | .485 | **.029** | .611 |
| **Age** | 58.6±4.89 | 77.3±3.38 | 61.0±4.82 | 76.2±3.92 | 69.2±9.76 | **.005** | **.008** | - |
| Range | 51.0 – 69.0 | 70.3 – 85.0 | 50.9 – 69.4<sup>a</sup> | 70.1 – 85.1 | 50.9 – 85.1 |  |  |  |
| **Education (years)** | 13.2±3.12 | 12.4±3.74 | 14.1±3.33 | 12.5±4.79 | 12.8±3.86 | .116 | .816 | .052 |
| Missing | 2 (1.1) | 0 (0.0) | 1 (2.4) | 6 (3.9) | 9 (1.7) |  |  |  |
| **APOE-ε4 allele +** | 85 (45.2) | 29 (19.2) | 25 (61.0) | 114 (74.0) | 253 (47.4) | .066 | **<.001** | .116 |
| **CSF Aβ42/40 +<sup>b</sup>** | 0 (0.0) | 0 (0.0) | 41 (100) | 154 (100) | 195 (36.5) | - | - | - |

Continuous variables are displayed as mean±SD. Categorical variables are displayed as n (%). <sup>a</sup> individuals who reported an age-of-onset under 65 were included in the EOAD group. <sup>b</sup> Aβ positivity: <.08 on CSF Aβ42/40 ratio.
Abbreviations: Aβ=amyloid-beta; AD=Alzheimer's disease; CU=cognitively unimpaired; CSF=cerebrospinal fluid; EOAD=early-onset cognitive impairment, LOAD=late-onset cognitive impairment; MCI=mild cognitive impairment; OCU=older cognitively unimpaired controls; SD=standard deviation; YCU=younger cognitively unimpaired controls.

### Amnestic EOAD shows medial temporal lobe subfield involvement

A statistically significant difference in mean value was found for all MTL regions of interest for both EOAD and LOAD compared to respective control groups (Fig. 1, supplementary results sTable 3). The biggest differences comparing EOAD with controls were observed in amygdala, BA35, and total hippocampus (z-score mean differences = 1.89, 1.70, - 1.68 respectively, all p<.001). The biggest differences comparing LOAD with controls were observed in entorhinal cortex, amygdala, and total hippocampus (mean differences = 1.59, 1.55, 1.55 respectively, all p<.001). These results indicate similar atrophy patterns across the medial temporal lobe (sFig. 11) between EOAD and LOAD, which was also confirmed by the lack of statistically significant interactions between age and diagnosis (sTable 4). The only exception was ERC where larger atrophy in LOAD appears and the interaction between age group and diagnosis was significant (sTable 4). These results contrast with previous reports which suggested limited involvement of the MTL in EOAD (see sTable 3). Including age as covariate did not change these results (see sTable 3).

**Figure 1.**
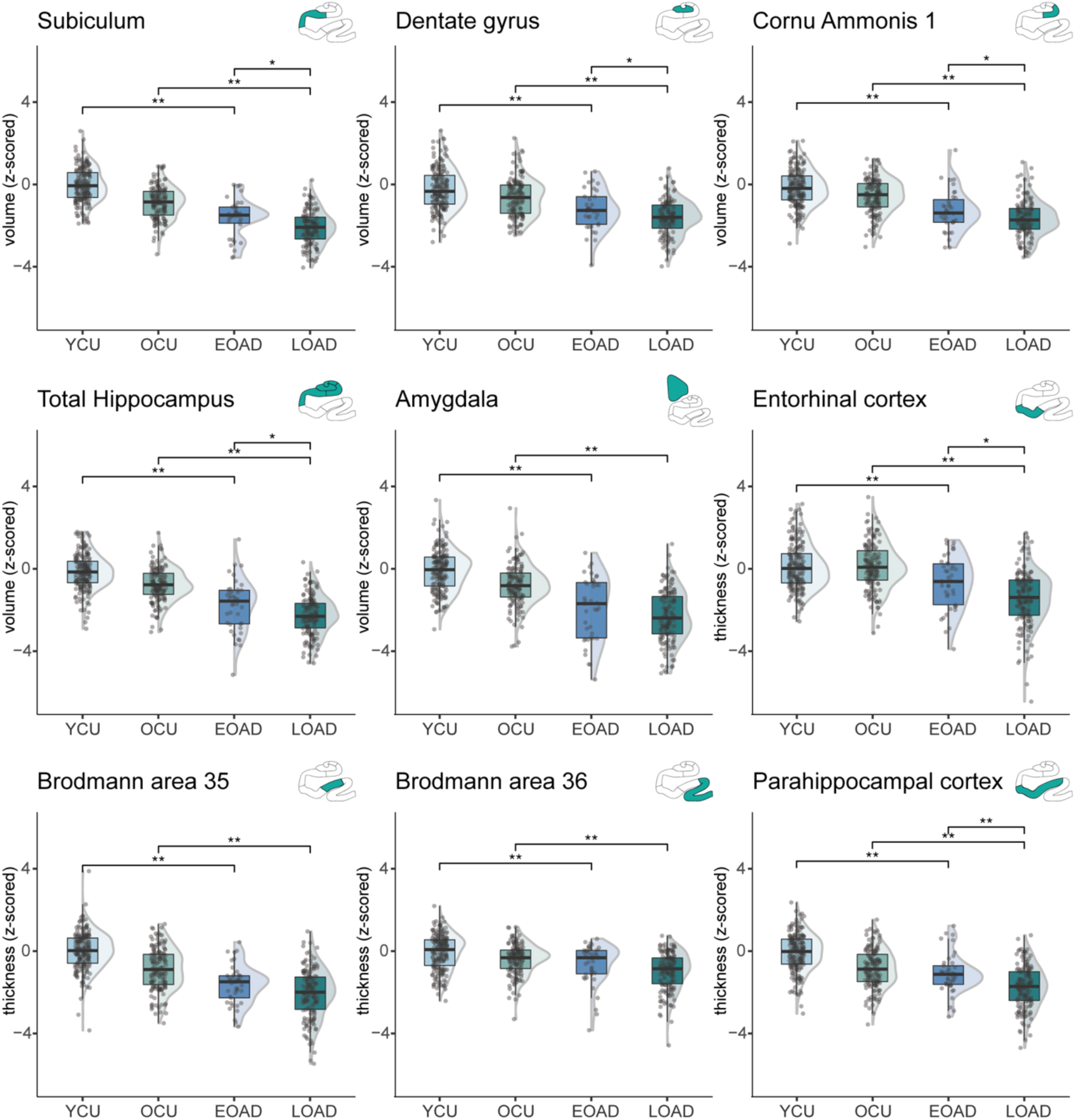
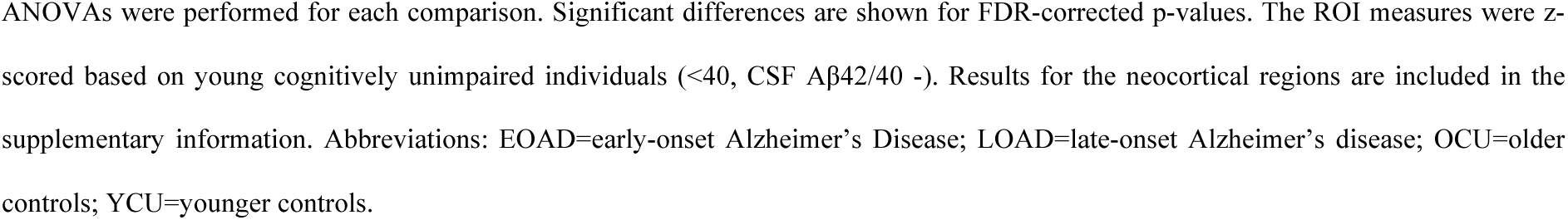
EOAD and LOAD group differences in medial temporal lobe subfield volume/thickness.

Focusing on the differences between EOAD and LOAD, significantly lower volume or thickness was found in LOAD compared to EOAD in five regions: subiculum (mean difference=0.50, *p_FDR_*=0.004), dentate gyrus (mean difference=0.38, *p_FDR_*=0.042), Cornu Ammonis 1 (mean difference=0.50, *p_FDR_*=0.042), entorhinal (mean difference=0.78, *p_FDR_*=0.003), and parahippocampal cortex (mean difference=0.68, *p_FDR_*<0.001). Also, total hippocampal volume differed between EOAD and LOAD (mean difference=0.50, *p_FDR_*=0.011, Fig. 1, sTable 3).

### Further characterization of amnestic EOAD and LOAD

#### Neocortical thickness differences in amnestic EOAD vs. LOAD for frontal and lateral temporal cortices

As additional analyses, potential differences in thickness of neocortical regions in EOAD and LOAD were investigated. When comparing AD groups with their respective controls, a statistically significant difference was found for all neocortical regions of interest (sFig. 12A). The pattern of atrophy between EOAD and LOAD compared with respective controls was similar for all regions except for lateral and medial parietal cortices, for which the interaction between age group and diagnosis was also significant, indicating more prominent atrophy in the EOAD group (sFig. 12B). Additionally, significantly lower lateral temporal and frontal thickness was found in LO-compared to EOAD (*p_FDR_*=0.031, 95%-C.I.=[-0.728, -0.051] and *p_FDR_*=0.014, 95%-C.I.=[-0.841, -0.116] respectively; sFig. 12).

Lastly, comparisons of thickness in the LEADS signature were performed. Both EOAD and LOAD showed significantly thinner thickness compared to controls but no differences between EOAD and LOAD were observed (see sFig. 13).

#### Amnestic EOAD shows a similar AD- and co-pathology burden as amnestic LOAD compared to controls

In a next step, we investigated potential differences in amnestic EOAD vs. LOAD with regards to common pathologies often accumulating in and related to MTL atrophy. Comparing AD groups with respective controls, a statistically significant difference in mean value was found for most AD pathologies and co-pathologies, indicating significantly higher pathology burden in the AD groups (MTL tau-PET SUVR, aHC/PHC ratio as TDP-43 proxy, CSF Aβ42/Aβ40 ratio; Fig. 2, supplementary material sTable 5). Only the total volume of WMH did not differ significantly between LOAD and controls (*p_FDR_*=.085). The results remained consistent when including age as covariate, except that a significant difference between LOAD and controls was found for WMH (*p_FDR_*=.033, see supplementary results sTable 6).

**Figure 2.**
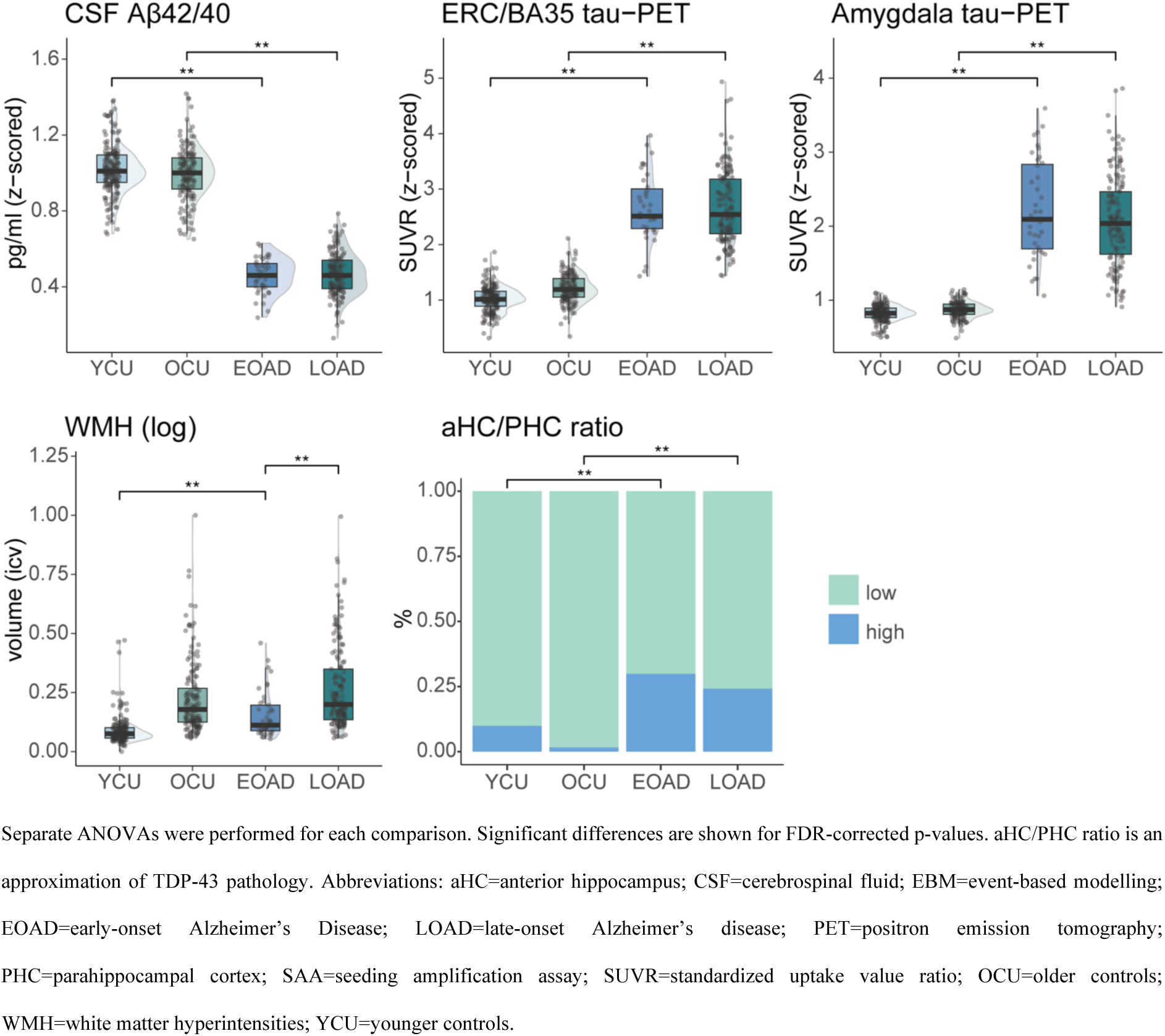
EOAD and LOAD group differences in AD pathologies and co-pathologies.

Focusing on the differences between EOAD and LOAD, we found a statistically significant higher mean value for WMH in LOAD compared to EOAD (*p_FDR_*<0.001, 95%-C.I.=[0.056, 0.170]; Fig. 2; see supplementary sTable 5 and sFig. 14 for results using a dichotomized white matter hyperintensity measure). No differences in biomarkers of AD (MTL tau-PET and CSF Aβ42/Aβ40 ratio) were observed between EOAD and LOAD (Fig. 2). Additionally, no differences between EOAD and LOAD were observed in the proportion of positivity for MRI-based proxy of TDP-43 pathology (Fig. 2, sTable 5). Results of these group comparisons did not change when accounting for CSF Aβ42/Aβ40 ratio in the models.

Comparisons of tau-PET uptake in all four neocortical composite regions and the LEADS signature show higher uptake in AD groups compared to controls and EOAD showed a significantly higher tau-PET uptake in these neocortical composite regions compared to LOAD (see sTable 5, sFig. 13 and sFig. 15).

#### Association between AD- and co-pathologies and atrophy in amnestic EOAD

In order to explore potential associations between AD- and co-pathologies and the structural measures, we focused only on the regions of interest which showed significant differences between EOAD and LOAD (total hippocampus (including subiculum, dentate gyrus, and cornu ammonis 1), entorhinal, parahippocampal; see sFig. 16).

Only the proxy of the presence of TDP-43 pathology was significantly associated with smaller total hippocampal volumes (std. *β*=-.63, *p_FDR_*<0.001). However, this association may be due to the definition of the measure considering the anterior hippocampus constitutes a large proportion of total hippocampal volume.

#### Cognitive performance in amnestic EOAD

Exploring group differences in cognitive performance, worse performance of the AD groups compared to respective controls was observed for all cognitive measures, while lower verbal fluency and naming abilities in LOAD compared to EOAD were observed (see sTable 7). No significant associations between MTL atrophy and performance on cognitive domains dependent on the MTL (episodic memory, naming, semantic fluency) were found for the amnestic EOAD group (see sFig. 17).

#### Comparison between amnestic and non-amnestic EOAD and LOAD

Demographic information on the non-amnestic AD (naEOAD: n=7; naLOAD: n=16) are provided in the supplementary material (sTable 8). Both amnestic AD groups showed lower MTL, but not neocortical, volume/thickness compared to non-amnestic AD (see sTable 9, sFig. 18-19). Subiculum volume and BA35 thickness were significantly smaller in amnestic vs. non-amnestic EOAD (see sTable 9, sFig. 18). The amnestic, compared to the non-amnestic AD groups showed higher amygdala tau-PET uptake. Non-amnestic LOAD showed larger WMH volumes compared to amnestic LOAD (see sTable 10, sFig. 20).

## Discussion

The major aim of this cross-sectional study was to investigate if the MTL is affected in amnestic EOAD by comparing this group to amnestic LOAD atrophy patterns and respective controls in fine-grained MTL subregions from a highly characterized cohort and using a new reliable automated whole amygdala segmentation. In contrast with previous reports (8–10), amnestic EOAD, as well as LOAD, showed significantly smaller volumes of MTL regions compared to controls. LOAD, compared to EOAD, was found to have smaller volumes/thickness in the MTL only for hippocampus, entorhinal, parahippocampal cortex, and in the neocortical regions in lateral temporal and frontal cortex. To further characterize the AD groups, we focused on biomarkers of AD and non-AD pathologies that often affect the MTL. The EOAD grouped showed higher neocortical tau-PET uptake but lower WMH burden, compared to LOAD. However, no differences were observed for our proxy of TDP-43. Lastly, the proxy of TDP-43 positivity was associated with smaller hippocampal volumes indicating a potential involvement in driving atrophy in this region.

Our results show that the MTL is affected in amnestic AD, irrespective of age. This may seem in contrast with previous reports showing evidence of relative sparing of the MTL in EOAD (1,51,52). However, since prior studies commonly grouped all EOAD subtypes together with, for example, with PCA, PPA, or non-amnestic EOAD, except e.g. (53), it is possible that MTL atrophy in these studies was concealed by other phenotypes. The importance of the MTL in memory function (13), suggests that an amnestic type of AD should be associated with MTL atrophy, regardless of age of onset, a notion that is supported by our findings.

Even though we observed lower MTL thickness in amnestic EOAD when comparing with controls, LOAD still shows more atrophy within the MTL (e.g., lower thickness in entorhinal compared to EOAD). This may be due to several reasons. First, there may be non-specific aging effects on these cortical structures leading to more atrophy in the older patient group. Second, for some individuals, pathologies may have a longer duration of accumulation in these regions, potentially exerting an effect on structure for a longer duration resulting in more atrophy. Previous reports of increased parietal atrophy in EOAD (1) were supported in our amnestic EOAD sample, given the significant interaction between age and diagnosis for parietal regions, indicating more prominent atrophy in the EOAD group than in LOAD. Additionally, we did observe higher levels of tau-PET uptake in parietal regions in the EOAD group, which may potentially contribute to the more pronounced atrophy in this region.

In comparison to respective controls, the amnestic AD groups show similar significant increased frequency or severity in the investigated co-pathologies. The only exception was observed for WMH which were increased in EOAD, but not in LOAD, where the results were more inconsistent. The fact that the EOAD group shows a similar level of co-pathologies as LOAD may be due to faster accumulation of pathologies, such as tau, but could also reflect a lack of resilience to pathologies. The mechanisms behind the presence of these co-pathologies for EOAD despite younger age remains to be elucidated.

It is of interest that no differences between EOAD and LOAD were found for a common co-pathology, the proxy of TDP-43 pathology. Previously, it has been reported significantly less TDP-43 proteinopathy in EOAD compared to LOAD (5). This was not replicated in the present study using a proxy of TDP-43 based on the observed anterior to posterior gradient of TDP-43 occurrences in the MTL (41). It is possible that the proxy, established in an autopsy cohort, does not replicate to our cohort, even though a similar cut-off was found when replicating it in our cohort (693 vs. 645) using Gaussian mixture modeling without postmortem validation. The fact that no difference between AD groups was observed could, however, also be due to a smaller sample size compared to what the study by Spina and colleagues (5) included and the indirect nature of our measure for presence of TDP-43. Nevertheless, we did find that our measure of TDP-43 positivity was associated with lower hippocampal volume in the amnestic EOAD group. Lastly, previous studies have reported a higher burden of AD pathology in amnestic EOAD compared to LOAD (2,5,54). We found that amnestic EOAD shows more neocortical tau pathology while presenting similar levels of MTL tau to LOAD. Our results are, thus, in line with the notion of EOAD showing a more aggressive disease progression with faster cognitive decline and accumulation of pathology (1) and previous observations of higher levels of tau accumulation in younger individuals (55). The null results regarding our analyses associating co-pathologies with MTL structural measures in EOAD are likely due to limited power.

### Strengths and Limitations

Strengths of the current study include the fine-grained investigation of MTL subfields, the use of a highly characterized cohort with various biomarkers of (co-)pathologies available, and the focus on amnestic EOAD as a separate group. Additionally, a new reliable automated segmentation for the whole amygdala is presented. However, the study also presents some limitations. First, the sample size of the EOAD group is relatively small. While this corresponds to the lower proportion of EOAD in the general population (56), it results in lower statistical power. Thus, future studies should investigate a larger sample of amnestic EOAD. Second, the cross-sectional nature of the study does not allow us to draw conclusions about potential more aggressive courses or larger atrophy rates between groups.

### Conclusions

In summary, we found a largely similar MTL atrophy pattern in amnestic EOAD compared to LOAD. Interestingly, besides lower white matter hyperintensity volumes and higher neocortical tau PET in EOAD compared to LOAD, no differences in other AD- and co-pathologies, such as MTL tau-PET, and our proxy of TDP-43 were observed between EOAD and LOAD. These results suggests that the driving mechanisms of the amnestic symptoms in both groups might be largely similar and resulting in similar atrophy patterns within the MTL.

## Data availability

Pseudo-anonymized data from BioFINDER-2 will be shared on request from a qualified academic investigator for the sole purpose of replicating procedures and results presented in the article and as long as data transfer is in agreement with EU legislation on the general data protection regulation and decisions by the Swedish Ethical Review Authority and Region Skåne, which should be regulated in a material transfer agreement.

## Abbreviations

Aβ: Amyloid-beta
AD: Alzheimer’s disease
aHC: anterior Hippocampus
AMY: Amygdala
ANCOVA: Analysis of covariance
ASHS: Automated Segmentation of Hippocampal Subfields
BA35: Brodmann area 35 (≈transentorhinal cortex)
CA1: cornu ammonis 1
C.I.: confidence interval
CSF: cerebrospinal fluid
EOAD: early-onset Alzheimer’s Disease
ERC: entorhinal cortex
LOAD: larly-onset Alzheimer’s Disease
MTL: medial temporal lobe
NFTs: tau neurofibrillary tangles
PET: positron emission tomography
pHC: posterior Hippocampus
PHC: parahippocampal cortex
TDP-43: transactive response DNA binding protein 43
WMH: white matter hyperintensities

## Supporting information

Supplementary Results

## Acknowledgements

We would like to acknowledge all the BioFINDER team members as well as participants in the study and their family members for their dedication. This study was supported by MultiPark - A Strategic Research Area at Lund University.

## Author information

Nicola Spotorno and Laura E.M. Wisse share co-last/equal authorship.

## Ethics declaration

### Ethics approval and consent to participate

The study was approved by the Regionala Etiksprövningsnämnden i Lund (regional ethical review board in Lund), Sweden, and all study participants provided written informed consent in accordance with the Declaration of Helsinki.

## Supplementary information

Supplementary material is available: supplementary methods and supplementary results.

