## Supplementary Methods for "Medial temporal lobe atrophy patterns in early- versus late-onset amnestic Alzheimer’s disease"

### Supplementary Material Methods
### Atrophy patterns and co-pathologies in early- versus late-onset amnestic Alzheimer's disease

### Amygdala segmentation protocol

Segmentation of the human amygdala was informed by atlases from Ding et al. (2018) and Mai, Paxinos, & Voss (2008). Segmentations were done on T1-weighted images with voxel size of 0.5x0.5x1.0 mm using ITK-SNAP (Yushkevich et al., 2006). Tracing was prepared in the coronal plane while the sagittal and axial plane were used to inspect and edit the segmentation. However, in case of conflicting information between the three planes, preference was always given to the coronal plane. The amygdala was segmented as a whole, and nuclei were not specified as subfields. The amygdala is traced in a posterior to anterior direction, starting with the most posterior slice of the amygdala because the borders of the amygdala can more easily be identified in posterior sections. The aim of the protocol is to only include amygdala tissue. However, as not all border of the amygdala can be clearly identified on MRI, it is possible that in some instances non-amygdala tissue will be included or that small portions of the amygdala will not be included in the final segmentation. More details are provided below.

The protocol is shown with one example case, but it is important to note that individual anatomical differences exist, so the shape of the amygdala may look somewhat different than in the shown example case. Furthermore, the protocol can be used to segment amygdalae in both hemispheres.

#### Posterior Portion of the Amygdala

##### *Most posterior slice*

The most posterior slice of the amygdala is defined as the first slice where hippocampal amygdaloid transition area (HATA) and the amygdala are not connected anymore (compare Fig. 1.2 and 1.3; see Fig. 2). In case of an ambiguous slice where the HATA and amygdala may be connected, the most posterior slice with a definite connection is chosen to determine the most posterior slice. This slice was chosen as the most posterior slice since it is consistently identifiable. However, a very small portion of the posterior tip of the amygdala may be excluded due to this rule. On this most posterior slice, the superior border is the white matter and the optic tract; the inferior border is the cerebrospinal fluid (CSF) in the temporal horn of the lateral ventricle (TLV); the medial border is CSF; the lateral border is the white matter of the temporal stem and the amygdalostriatal transition area (AStr). The inferolateral border should be symmetrical to the inferomedial side to exclude most of the tail of caudate (CAT). Fig. 2 (see arrows on bottom rows) shows that the inferolateral border does not extend further inferiorly than the inferomedial border.

Working in an anterior direction, in the next slice there is a definite connection between the HATA and the amygdala show a definite connection (Fig. 1.2, see Fig. 2). The superior, lateral, and inferior borders do not change, while the medial border is now CSF and the most superior portion of the HATA, which is not segmented as amygdala. The border to the HATA is determined by finding the shortest connection between the superomedial CSF and superior CSF (Fig. 3).

Continue moving anteriorly segmenting with the same borders as before (Fig. 1.3-1.5).

#### Mid-portion of the Amygdala

The mid-portion starts after a few slices (in the anterior direction) when a superior bulge relative to the endorhinal sulcus appears (starting Fig. 1.6). At this point the superiomedial border is not drawn straight but with a slight angle to the superior white matter. The inferiomedial border of the amygdala to the HATA is the semiannular sulcus (sas) (Fig.1.6). From the semiannular sulcus (noted in Fig. 1.7) the border is traced superolateral towards the fundus of the endorhinal sulcus and then continues in a slight angle to the superior white matter (see Fig. 4). The inferior border continues to be CSF and the hippocampus (HC).

Laterally, connect the superior white matter (i.e., the superior border) to the inferior CSF of the TLV (i.e., the inferior border) (see Fig. 4). To draw this line, use the gray/white matter border. In some cases, some grey matter may be lateral beyond the direct connection between superior white matter and inferior CSF (see Fig. 4), which would result in a biologically implausible bulge on the superolateral portion of the amygdala. If this is the case, this extra portion should not be included. In such cases, superimposing the border from the immediate posterior slice will additionally help to determine whether this portion should be excluded. Hence, it helps to move between slices when tracing these borders. This border is drawn so the AStr and putamen/peduncle of lentiform (PUV/PedL) are excluded from the segmentation. Continue moving anteriorly segmenting with the same borders (Fig. 1.7).

When moving more anteriorly, the inferior border of the amygdala is the hippocampus and inferomedial the entorhinal cortex (ERC; Fig. 1.8). The border to the entorhinal cortex continues to be based on the semiannular sulcus. Since the border to the hippocampus is sometimes easier to identify in sagittal plane, the border to the hippocampus should be inspected in this plane to ensure correct segmentation.

Segmentation continues anteriorly with the same borders (Fig. 1.9-1.10).

*Last slice of the HC*

Segmentation continues as on the previous slices. Inferior, the TLV may start to decrease in some cases (e.g., compare Fig. 1.10 and 1.11). Therefore, the white matter of the parahippocampal gyrus will slowly become the new inferior border (see consecutive slices). If the TLV remains larger, continue segmenting the inferior-lateral border in the same fashion as on previous slices.

#### Anterior Portion of the Amygdala

*First slice anterior to the HC*

The superior border (white matter) continues to be traced from the sas (medial border) in a superolateral direction towards the fundus of the endorhinal sulcus and then continues in a slight angle (counterclockwise) to the superior white matter (Fig. 1.11). The medial border continues to be the sas. The inferomedial border of the amygdala is the parahippocampal white matter, which is segmented by following the white matter border from the inferior border (the parahippocampal white matter) to the sas (Fig. 1.12 and see Fig. 5 as example). The lateral and superolateral border continues to be the white matter of the parahippocampal gyrus. Inferolateral, CSF continues to be the border until the white matter of the parahippocampal gyrus becomes the border (see next slice).

Segmentation continues as on the previous slice. Inferiorly, the white matter of the parahippocampal gyrus becomes the border (Fig. 1.12). In the anterior portion of the amygdala, small parts of the preamygdalar claustrum (PACI) may be included in the segmentation, if PACI cannot be distinguished as a separate nucleus but rather seems fused with the amygdala (Fig. 6). As much of the PACI as possible should be excluded from the amygdala.

*First slice excluding the cortex*

According to the atlases (Mai, Paxinos, & Voss, 2008; Ding et al., 2018), the cortical nuclei only appear approximately 2.9 mm after the most anterior slice of the amygdala. Therefore, the three most anterior slices (3 mm) are segmented excluding the periamygdaloid cortex (PAM) and piriform cortex, temporal pole (PirT), which are not part of the amygdala (Fig. 1.13). In order to establish the last three anterior slices, one should determine the most anterior slice first (see next section).

Once the three most anterior slices are determined, segment the first slice (third most anterior slice) excluding the cortex in the following way: If there is a thin white matter layer between the cortex and the amygdala, that should be the superior and medial borders. If the white matter layer is not visible, the thickness of the cortex adjacent to the amygdala should be of the same thickness as the entorhinal cortex inferomedial to the amygdala, under the assumption that the thickness of the cortex is roughly the same. The inferior, inferomedial, and inferolateral borders continue to be the white matter of the parahippocampal gyrus and the grey matter of the entorhinal cortex.

Continue segmenting the next amygdala slice as on the previous slice (Fig., 1.14) ensuring that the amygdala is getting smaller with each slice moving anterior.

*Most anterior slice*

The most anterior slice is determined by visually assessing the coronal slices while again supporting the decision with the sagittal view (Fig. 1.15). Instead of the amygdala, white matter of the parahippocampal gyrus will occur in its place (Fig. 1.16), which is the first slice anterior of the amygdala that should not be segmented anymore. If there is a slice where grey matter of the amygdala is seen, but on the slice immediately anterior, the white matter of the parahippocampal gyrus occurs in its place, this definite slice with amygdala grey matter should be segmented as the most anterior slice. It can occur that there is a slice where there is mixed white and grey matter signal in the area where the amygdala was located on previous slices, but it is unclear whether it is amygdala or partial volume effects, this slice should be segmented as the most anterior slice of the amygdala. This rule should be followed for one slice only. If there are two slices where this occurs, only the one immediately anterior to the last definite slice should be segmented as amygdala. Note that there may be grey matter on a slice, but in a different location and is outside the area where the amygdala was previously located. This should not be segmented as amygdala and is not the most anterior slice. See Fig. 7 for examples of the most anterior slice.

The borders are the same as on the previous slice.

#### Finishing the segmentation

After segmenting the amygdala according to the protocol, it is helpful to go through the sagittal and axial planes to double-check the segmentation. First, it should be noted whether the border to the HC is correctly segmented (see Fig. 8). Second, all other borders should be checked and made smoother if possible. Third, after going through sagittal and axial view and potentially adjusting the segmentation, the coronal plane should be inspected again to ensure that the amygdala is correctly segmented according to the protocol. This process can be repeated if needed. It can be helpful to only make small adjustments in the sagittal and axial planes, to prevent deviations from the protocol in the coronal plane. Otherwise, it can be helpful to only make bigger changes in the axial and sagittal planes while inspecting all three planes at the same time.

#### Differences in amygdala segmentations

Fig. 10 exemplifies the segmentation of a different amygdala in the right hemisphere and its segmentation. Note that the overall shape is similar for both examples, while differences, such as the number of slices (14 slices vs. 16 slices), occur. Additionally, the endorhinal sulcus in Fig. 10 is much wider.

The semiannular sulcus may be challenging to identify on each slice and may differ per subject/hemisphere. Therefore, marking the sulcus where it can be clearly seen and extrapolating from these slices to slices on which this border is less clear may be helpful.

**Abbreviations**

| | |
|---|---|
| AMY | amygdala |
| Astr | amygdalostriatal transition area |
| CSF | cerebrospinal fluid |
| ERC | entorhinal cortex |
| es | endorhinal sulcus |
| HATA | hippocampal amygdaloid transition area |
| HC | hippocampus |
| opt | optic tract |
| PACI | preamygdalar claustrum |
| PAM | periamygdaloid cortex |
| PirT | piriform cortex, temporal pole |
| PUV/PedL | putamen/peduncle of lentiform |
| sas | semiannular sulcus |
| TLV | temporal horn of the lateral ventricle |

**Figure 1.** Example whole amygdala segmentation moving from a posterior (1.1) to an anterior direction. For each slice the raw MRI, an annotated version, and a segmented version is shown in a row.

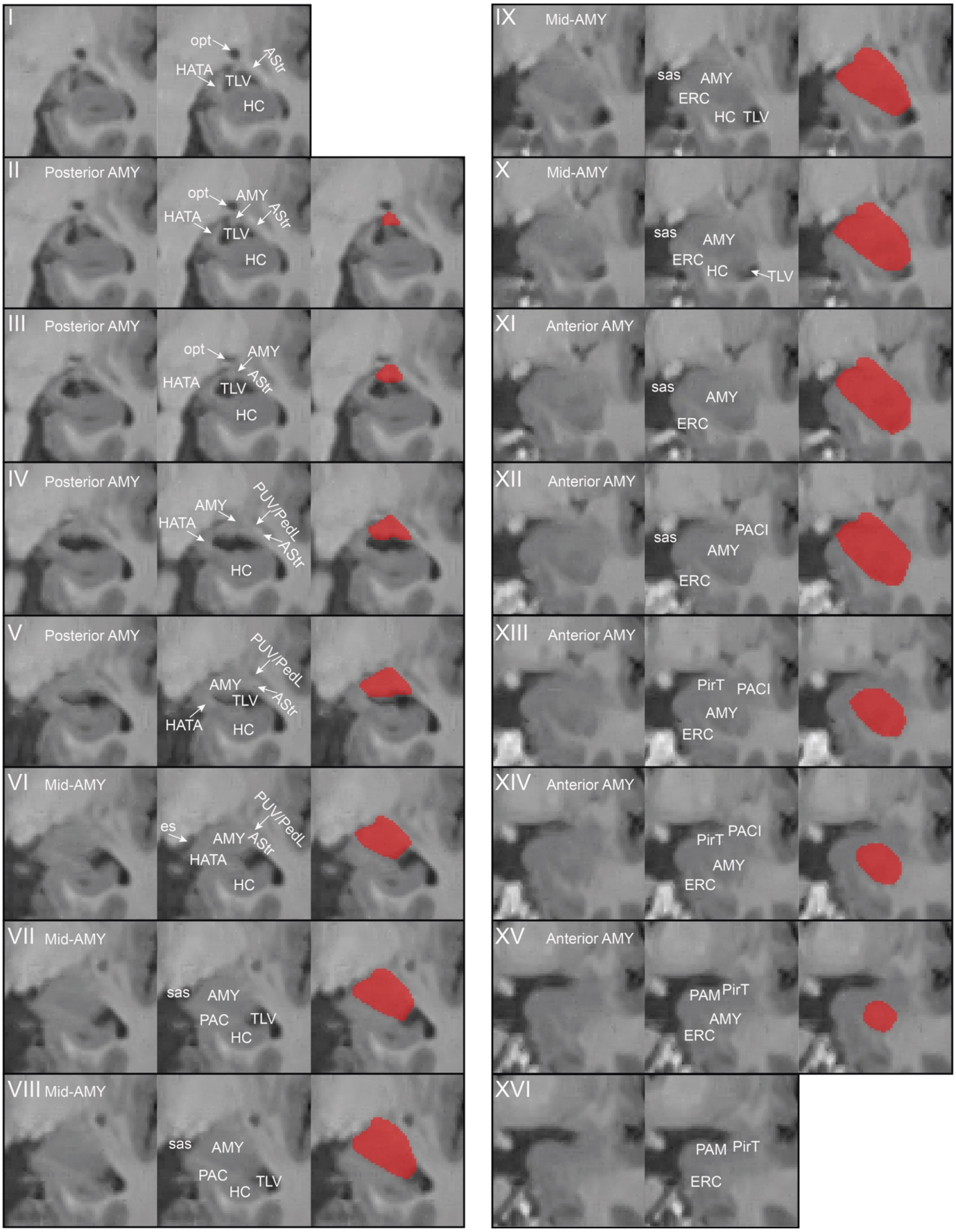

*Abbreviations*: AMY: amygdala; AStr: amygdalostriatal transition area; CSF: cerebrospinal fluid; ERC: entorhinal cortex; es: endorhinal sulcus; HATA: hippocampal amygdaloid transition area; HC: hippocampus; opt: optic tract; PACI: preamygdalar claustrum; PAM: periamygdaloid cortex; PirT: piriform cortex, temporal pole; PUV/PedL: putamen/peduncle of lentiform; sas: semiannular sulcus; TLV: temporal horn of the lateral ventricle.

**Figure 2.** Two examples of most posterior slice of the amygdala (A and B). Case A is a well-defined example where the connection between hippocampal amygdaloid transition area (HATA) and amygdala is clearly visible. Case B shows an ambiguous slice (B.2). Since this is unclear, the last slice where there is no ambiguous connection (B.3) is used to determine the most anterior slice.

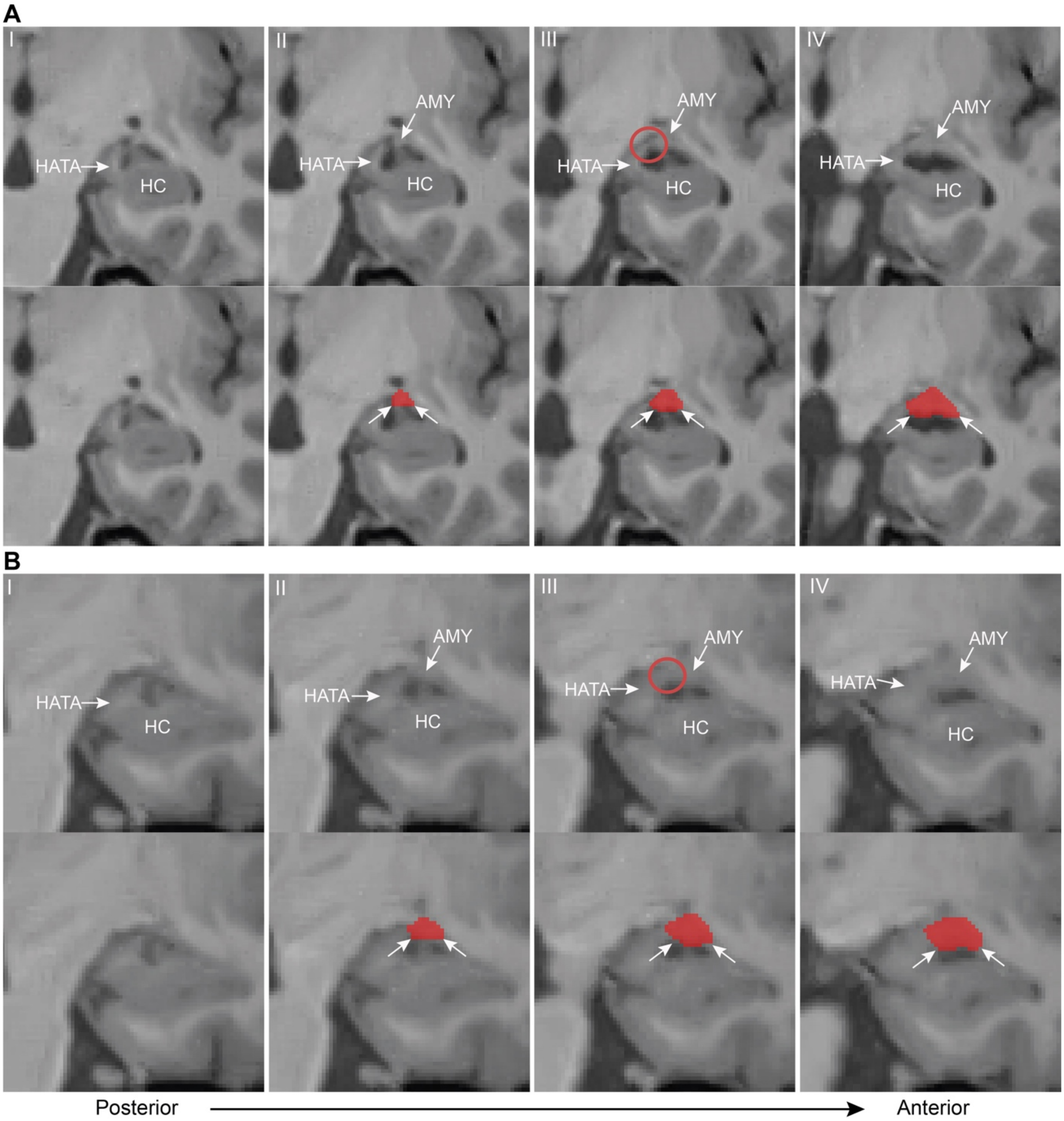

*Abbreviations*: AMY: amygdala; HATA: hippocampal amygdaloid transition area; HC: hippocampus.

**Figure 3.** The medial border of the posterior portion of the amygdala to the hippocampal amygdaloid transition area (HATA) if found by outlining the grey matter and finding the shortest connection between the superomedial cerebrospinal fluid (CSF) of the endorhinal sulcus (es) and superior CSF of the temporal horn of the lateral ventricle (TLV).

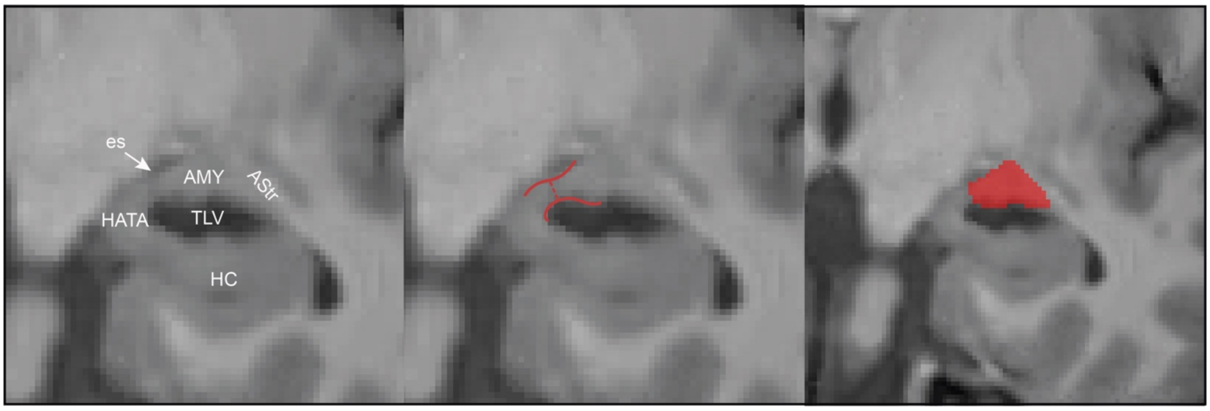

*Abbreviations:* AMY: amygdala; AStr: amygdalostriatal transition area; es: endorhinal sulcus; HATA: hippocampal amygdaloid transition area; HC: hippocampus; TLV: temporal horn of the lateral ventricle.

**Figure 4.** Lateral and superior border of the mid-portion of the amygdala showing differences in lateral borders and the superior white matter border. Panel A shows the biologically implausible bulge that should not be segmented as amygdala and in which case the white matter border is not followed (compared to panel B). This border is drawn so the putamen/peduncle of lentiform (PUV/PedL) is excluded (segmented in blue). Additionally, the medial white matter border of the amygdala which connects the semiannular sulcus and the temporal horn of the lateral ventricle is exemplified.

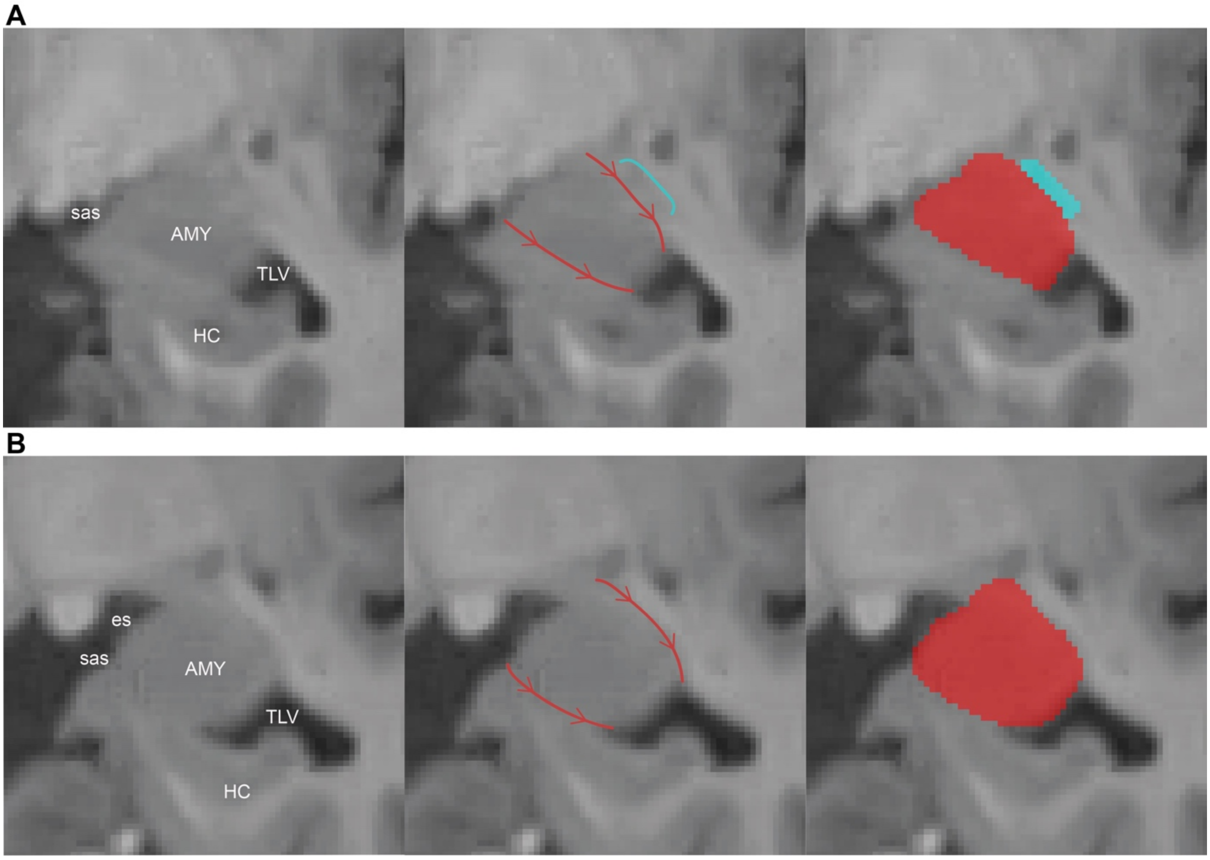

*Abbreviations:* AMY: amygdala; es: endorhinal sulcus; HC: hippocampus; sas: semiannular sulcus; TLV: temporal horn of the lateral ventricle.

**Figure 5.** The medial border of the anterior portion of the amygdala follows the white matter border from the semiannular sulcus to the parahippocampal white matter.

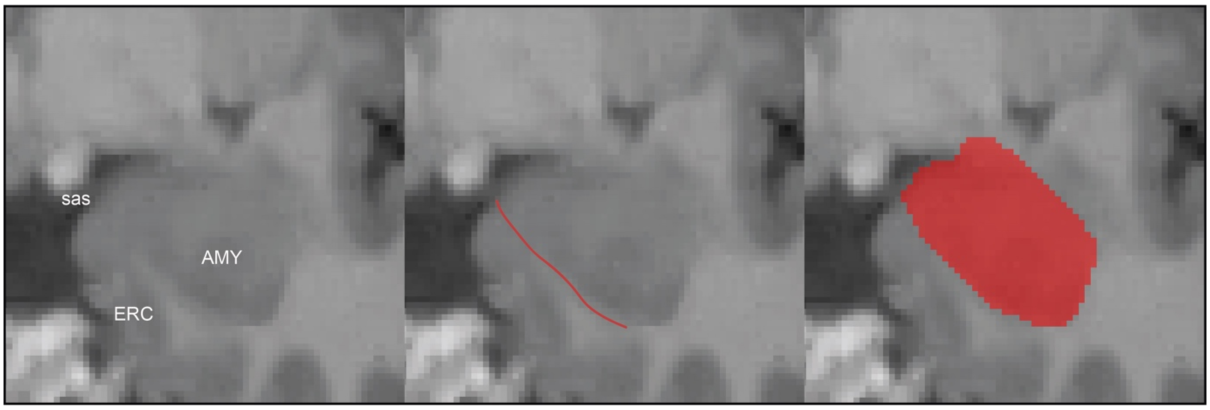

*Abbreviations:* AMY: amygdala; ES: endorhinal sulcu; sas: semiannular sulcus.

**Figure 6.** The preamygdalar claustrum (PACI), which is highlighted with the white circle is grey matter in the anterior portion of the amygdala which should be excluded from the amygdala segmentation.

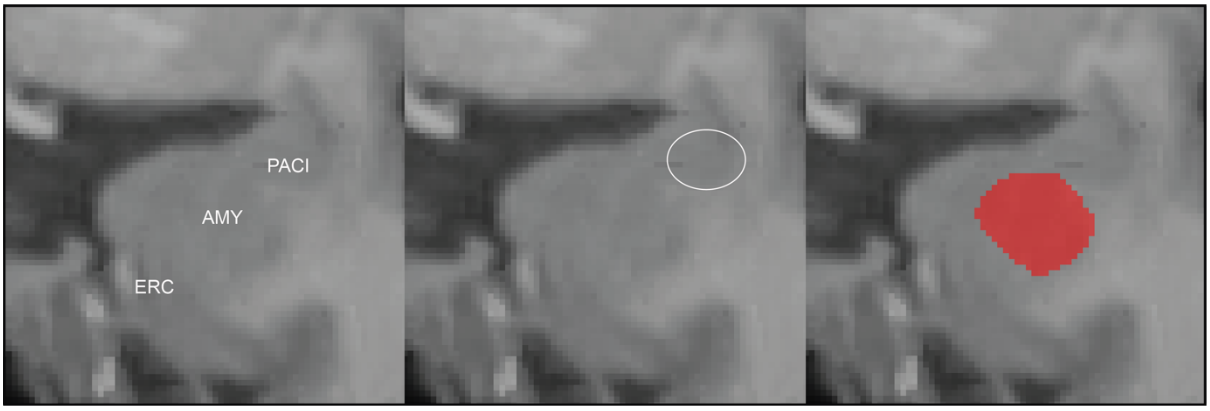

*Abbreviations:* AMY: amygdala; ERC: entorhinal cortex; PACI: preamygdalar claustrum.

**Figure 7.** The most anterior slice of the left (A) and right (B) amygdala is the last slice where the grey matter of the amygdala is seen and the white matter of the parahippocampal gyrus occurs in its place on the immediate anterior slice (purple line in A.3-4 and B.3-4). Note that the images do not capture the boundaries as well as moving between the slices in ITK-SNAP.

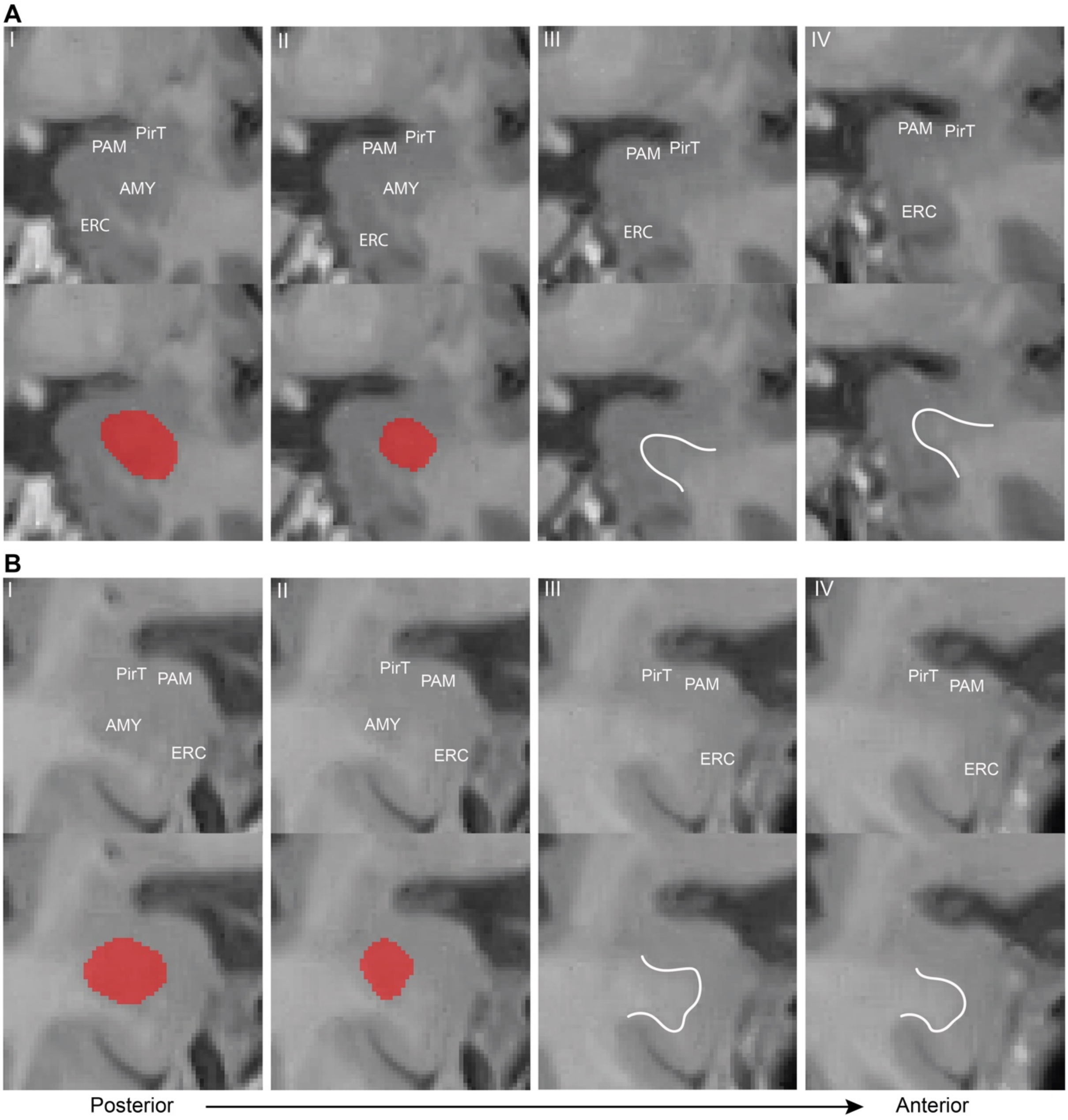

*Abbreviations:* AMY: amygdala; ERC: entorhinal cortex; PAM: periamygdaloid cortex; PirT: piriform cortex, temporal pole.

**Figure 8.** Sagittal view of the amygdala. In this view, the border to the hippocampus should be checked to ensure correct segmentation of the amygdala-hippocampus border.

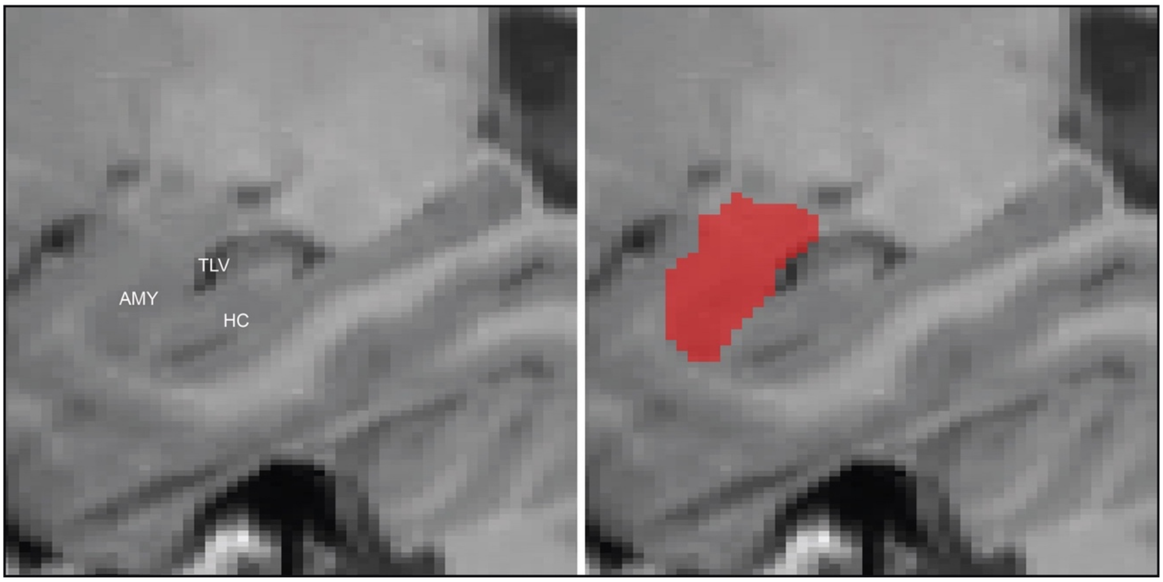

*Abbreviations:* AMY: amygdala; HC: hippocampus; TLV: temporal horn of the lateral ventricle.

**Figure 9.** Exemplary 3D view of a final amygdala segmentation.

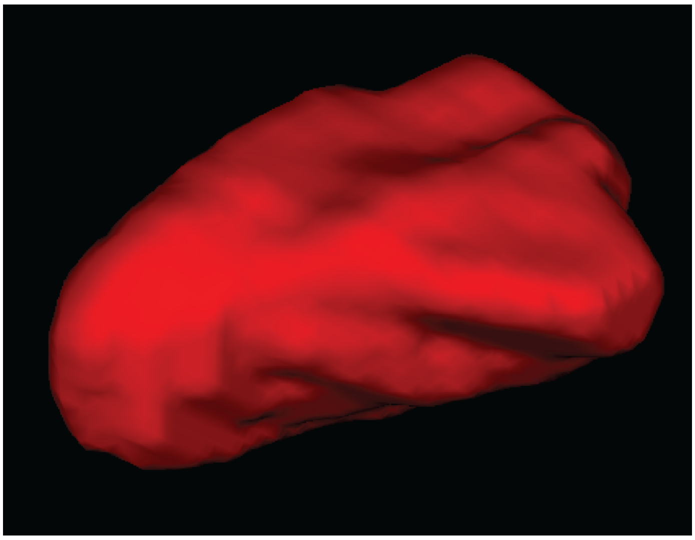

**Figure 10.** Second example segmentation of the whole amygdala showing the right hemisphere. The figure moves from a posterior (1.1) to an anterior direction. Each slice is shown as the raw MRI snapshot, an annotated version, and a segmented version in a row.

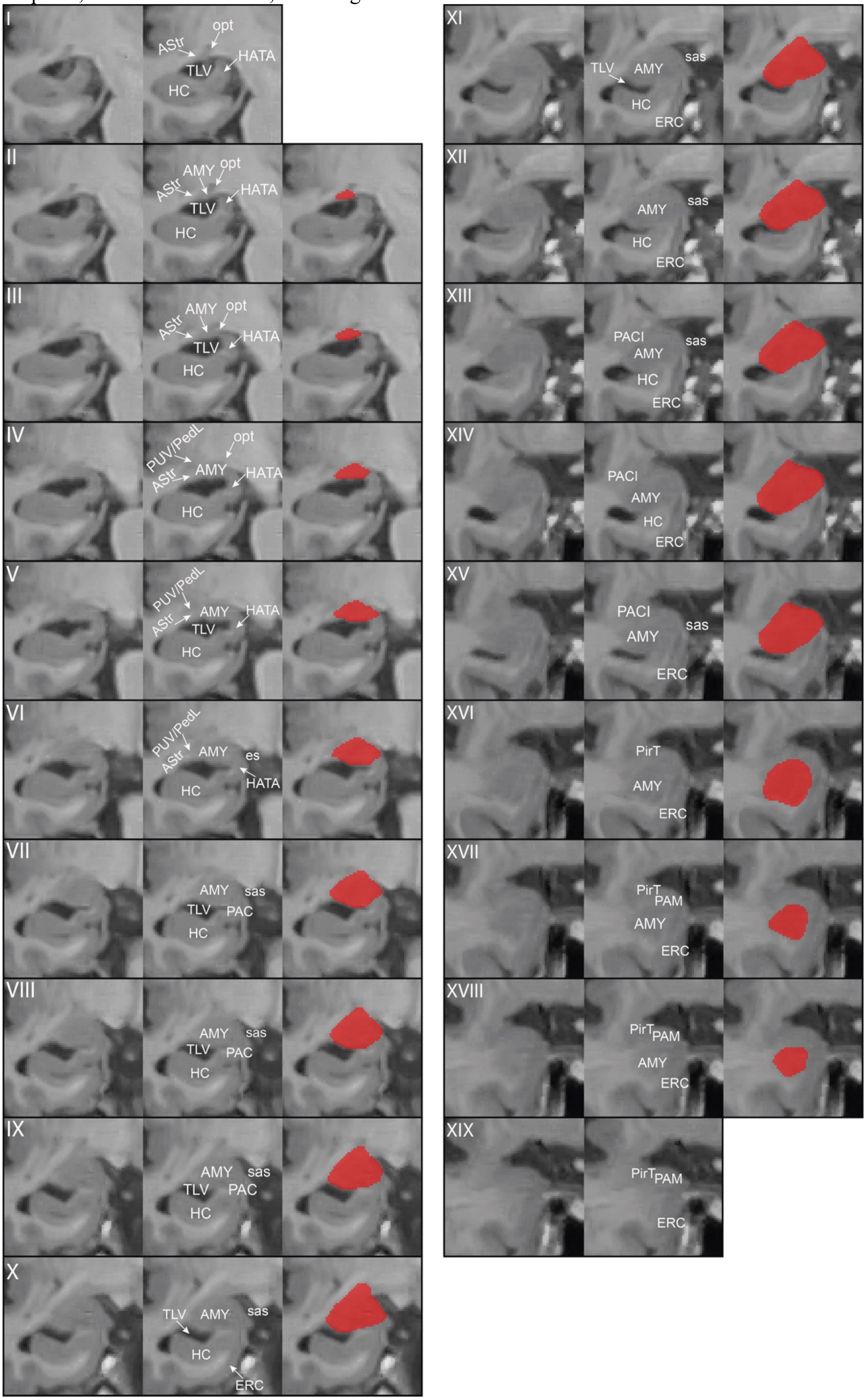

*Abbreviations*: AMY: amygdala; AStr: amygdalostriatal transition area; CSF: cerebrospinal fluid; ERC: entorhinal cortex; es: endorhinal sulcus; HATA: hippocampal amygdaloid transition area; HC: hippocampus; opt: optic tract; PACI: preamygdalar claustrum; PAM: periamygdaloid cortex; PirT: piriform cortex, temporal pole; PUV/PedL: putamen/peduncle of lentiform; sas: semiannular sulcus; TLV: temporal horn of the lateral ventricle.

### Reliability analyses

After protocol establishment a reliability test was performed. AW segmented the whole amygdala on both hemispheres in 15 subjects twice. Three weeks lied between the two segmentation instances. The segmentations were performed in a random order on both instances, which AW was blinded to. Additionally, AW was blinded to the diagnosis of the subjects (CU or MCI).

The results indicate high reliability for the amygdala both on intraclass correlation coefficient and the dice similarity index (sTable 1). The amygdala label was also added to the T1-ASHS atlas from Xie et a., (2019) and ASHS was retrained with the amygdala and all the subfields in the Xie et al., (2019) atlas. Cross-validation of manual versus automated segmentation was performed after training ASHS with the new atlas including the whole amygdala segmentation. The cross-validation dice similarity index of the amygdala was 0.88±0.03 (sTable 2). In comparison, the dice similarity index was also calculated for the anterior hippocampus resulting in 0.91±0.02; for the posterior hippocampus in 0.89±0.02; and for entorhinal cortex in 0.75±0.05 (see also Xie et al., 2019).

**sTable 1.** Intraclass correlation coefficient for the reliability of the whole amygdala segmentation protocol for the manual intra-rater reliability.

| | **ICC** | **lower 95%-CI** | **upper 95%-CI** |
|---|---|---|---|
| Left | .956 | .869 | .985 |
| Right | .961 | .884 | .987 |
| Left/Right | .985 | .955 | .995 |

**sTable 2.** Dice similarity index for the reliability of the whole amygdala segmentation protocol of the cross-validation using ASHS.

| | **Mean DSI** | **SD** | **Min** | **Max** |
|---|---|---|---|---|
| Left | .917 | .022 | .878 | .948 |
| Right | .910 | .027 | .863 | .941 |
| Left/Right | .914 | .015 | .888 | .940 |
