## Supplementary Results for "Medial temporal lobe atrophy patterns in early- versus late-onset amnestic Alzheimer’s disease"

### Supplementary Results
### Atrophy patterns and co-pathologies in early- versus late-onset amnestic Alzheimer's disease

### Amnestic EOAD shows medial temporal lobe subfield involvement

**sTable 3.** Comparison between groups on structural MRI measures.

| | YCU vs. EOAD | | | OCU vs. LOAD | | | EOAD vs. LOAD | |
|---|---|---|---|---|---|---|---|---|
| | diff | $p_{FDR}$ | $p_{FDR}$ age adjusted | diff | $p_{FDR}$ | $p_{FDR}$ age adjusted | diff | $p_{FDR}$ |
| **SUB** | 1.617 | **<.001** | **<.001** | 1.249 | **<.001** | **<.001** | 0.502 | **0.004** |
| **DG** | 1.043 | **<.001** | **<.001** | 0.993 | **<.001** | **<.001** | 0.380 | **0.042** |
| **CA1** | 1.124 | **<.001** | **<.001** | 1.122 | **<.001** | **<.001** | 0.387 | **0.042** |
| **ERC** | 0.808 | **<.001** | **<.001** | 1.598 | **<.001** | **<.001** | 0.779 | **0.003** |
| **BA35** | 1.698 | **<.001** | **<.001** | 1.196 | **<.001** | **<.001** | 0.409 | 0.058 |
| **BA36** | 0.608 | **<.001** | **<.001** | 0.601 | **<.001** | **<.001** | 0.337 | 0.060 |
| **PHC** | 1.035 | **<.001** | **<.001** | 0.913 | **<.001** | **<.001** | 0.677 | **<.001** |
| **Total HC** | 1.684 | **<.001** | **<.001** | 1.547 | **<.001** | **<.001** | 0.499 | **0.011** |
| **AMY** | 1.899 | **<.001** | **<.001** | 1.548 | **<.001** | **<.001** | 0.366 | 0.147 |
| **LT** | 1.314 | **<.001** | **<.001** | 1.036 | **<.001** | **<.001** | 0.381 | **0.031** |
| **LP** | 1.536 | **<.001** | **<.001** | 0.754 | **<.001** | **<.001** | -0.114 | 0.583 |
| **MP** | 1.439 | **<.001** | **<.001** | 0.849 | **<.001** | **<.001** | 0.049 | 0.741 |
| **FL** | 0.510 | **.001** | **.002** | 0.689 | **<.001** | **<.001** | 0.461 | **0.014** |
| **OL** | 0.371 | **.034** | **.037** | 0.276 | **.029** | **.009** | 0.157 | 0.438 |

Positive mean differences indicate higher values in the group listed first; negative mean differences indicate lower values in the group listed first. All analyses were adjusted for sex. All p-values are FDR adjusted.
Abbreviations: AMY=amygdala; BA=Brodmann area; CA1=cornu ammonis 1; DG=dentate gyrus; diff=mean difference; ERC=entorhinal cortex; EOAD=amnestic early-onset cognitive impairment; FL=frontal cortex; HC=hippocampus; LOAD=amnestic late-onset cognitive impairment; LT=lateral temporal; LP=lateral parietal; MP=medial parietal; naEOAD=non-amnestic early-onset AD; naLOAD=non-amnestic late-onset AD; OCU=older cognitively unimpaired controls; OL=occipital cortex; PHC=parahippocampal cortex; SUB=subiculum; YCU=younger cognitively unimpaired controls.

**sFigure 11.** Mean differences for the comparisons between EOAD with controls and LOAD with controls showing similar differences across regions.

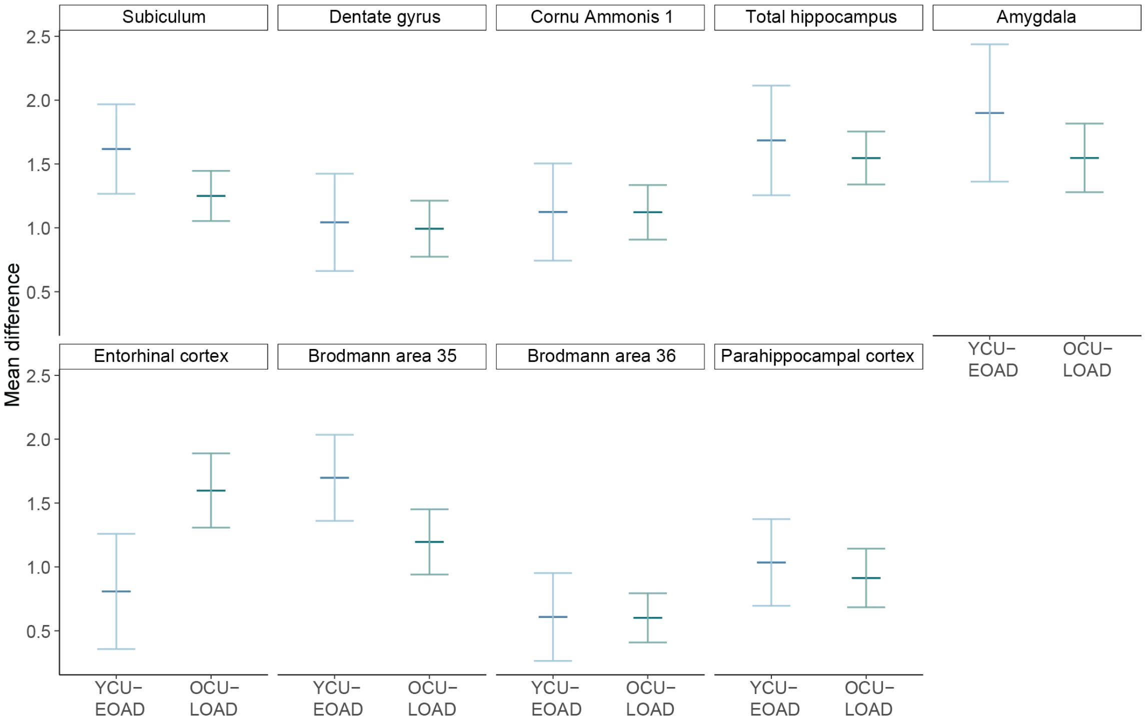

Mean differences in the groups are shown with a 95%-confidence interval. Abbreviations: AMY=amygdala; BA=Brodmann Area; CA1=Cornu Ammonis 1; DG=dentate gyrus; EOAD=early-onset Alzheimer's Disease; ERC=entorhinal cortex; FL=frontal cortexHC=total hippocampus; LOAD=late-onset Alzheimer's disease; LT=lateral temporal cortex; LP=lateral parietal cortex; MP=medial parietal cortex; OCU=older controls; OL=occipital cortex; PHC=parahippocampal cortex; SUB=subiculum; YCU=younger controls.

**sTable 4.** Results of the interaction analyses between age (young/old) and diagnosis (CU/AD) for all regions of interest.

| MTL subfields | Std. beta | 95%-CI | | $p_{FDR}$ |
|---|---|---|---|---|
| **SUB** | 0.134 | (-0.003 | 0.730) | 0.157 |
| **DG** | 0.017 | (-0.378 | 0.464) | 0.977 |
| **CA1** | -0.002 | (-0.400 | 0.389) | 0.977 |
| **ERC** | -0.262 | (-1,323 | -0.333) | **0.010** |
| **BA35** | 0.163 | (0.059 | 0.922) | 0.117 |
| **BA36** | -0.011 | (-0.371 | 0.323) | 0.977 |
| **PHC** | 0.049 | (-0.274 | 0.533) | 0.792 |
| **Total HC** | 0.046 | (-0.253 | 0.518) | 0.792 |
| **AMY** | 0.104 | (-0.138 | 0.824) | 0.363 |

The results show only the interaction term for each region of interest. The interaction between analyses were adjusted for sex. All p-values are FDR adjusted.

Abbreviations: AMY=amygdala; BA=Brodmann area; CA1=cornu ammonis 1; DG=dentate gyrus; diff=mean difference; ERC=entorhinal cortex; EOAD=amnestic early-onset cognitive impairment; HC=hippocampus; LOAD=amnestic late-onset cognitive impairmentOCU=older cognitively unimpaired controls; PHC=parahippocampal cortex; SUB=subiculum; YCU=younger cognitively unimpaired controls.

### Further characterization of amnestic EO- and LOAD

#### Neocortical thickness differences in EO- vs. LOAD

**sFigure 12.** EO- vs. LOAD group differences in neocortical volume/thickness.

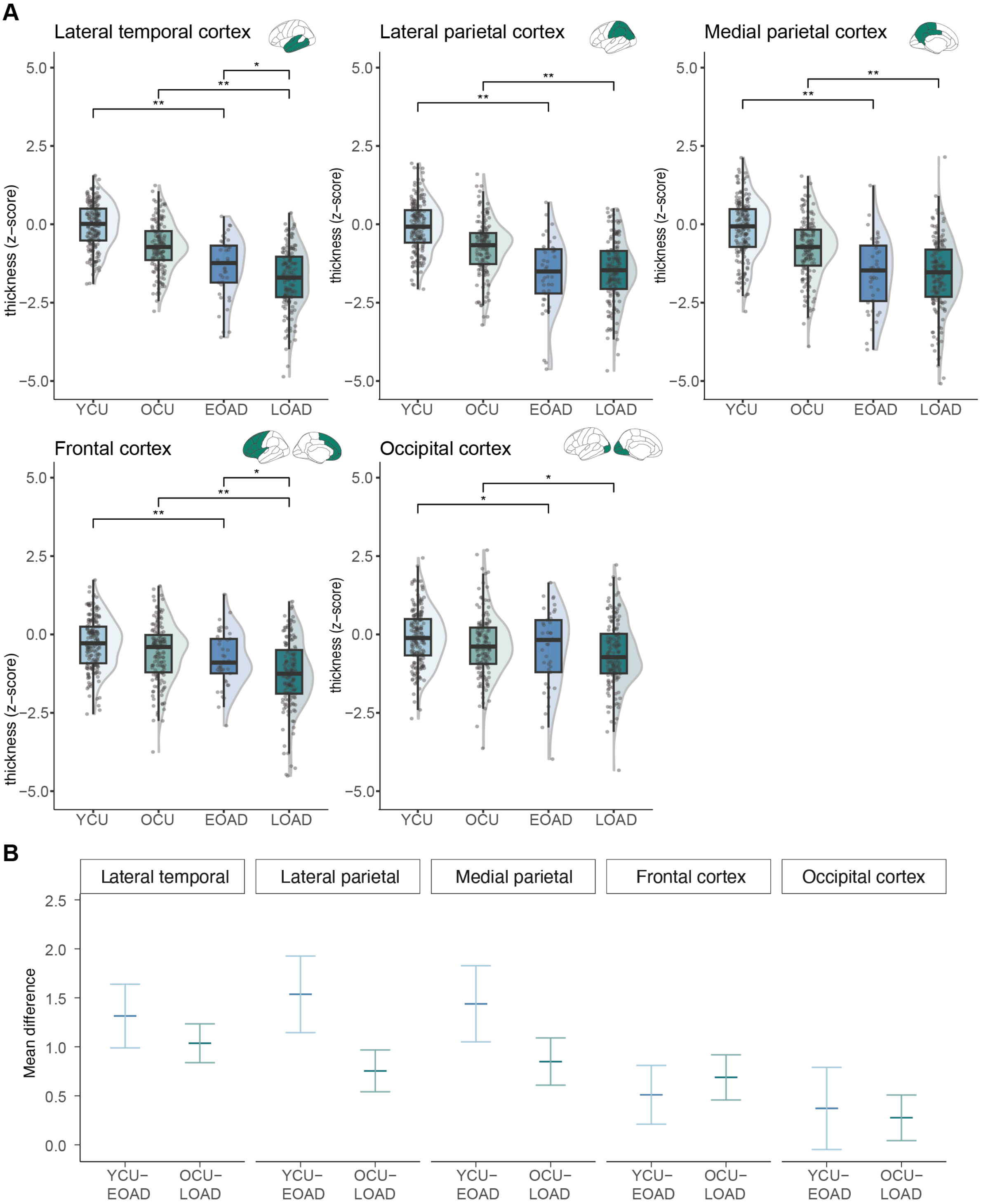

| Interaction | Lateral temporal | | Lateral parietal | | Medial parietal | | Frontal cortex | | Occipital cortex | |
|---|---|---|---|---|---|---|---|---|---|---|
| Std. beta | 0.121 | p=0.155 | 0.330 | **p<.001** | 0.230 | **p=0.010** | -0.070 | p=0.516 | 0.041 | p=0.653 |
| 95%-CI | -0.049 - 0.628 | | 0.436 - 1.172 | | 0.197 - 1.025 | | -0.536 - 0.220 | | -0.317 - 0.505 | |

**A** shows the group comparisons. ANOVAs were performed for each comparison. See sTable 6 for more information. **B** shows the mean differences of the comparisons of the AD groups with respective controls and the results of the interaction analysis (age*diagnosis) for the interaction term. Significant differences are shown for FDR-corrected p-values. Abbreviations: EOAD=early-onset Alzheimer's Disease; LOAD=late-onset Alzheimer's disease; OCU=older controls; YCU=younger controls.

#### LEADS signature thickness and tau-PET uptake group comparisons

**sFigure 13.** Group comparison for LEADS signature thickness and tau-PET uptake.

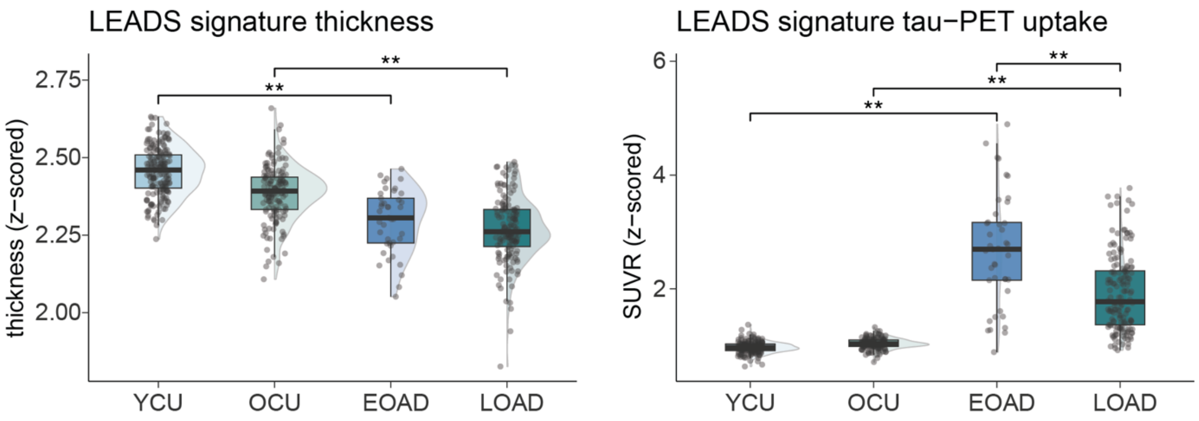

Significant differences are shown for FDR-corrected p-values for the exploratory analyses investigating the LEADS signature for thickness and tau-PET uptake. LEADS signature based on Touroutoglou et al. (2023).
Abbreviations: EOAD=early-onset Alzheimer's Disease; LOAD=late-onset Alzheimer's disease; LEADS=Longitudinal Early-Onset Alzheimer's Disease Study; PET=positron emission tomography; SUVR=standardized uptake value ratio; OCU=older controls; YCU=younger controls.

Both EO- and LOAD showed significantly thinner thickness compared to controls ($p$<0.001, 95%-C.I.=[-0.20, -0.12]; $p$<0.001, 95%-C.I.=[-0.14, -.08] respectively). No differences between EO- and LOAD were observed (p=.351, 95%-C.I.=[0.02, 0.35]).

Both EO- and LOAD showed significantly higher tau-PET uptake compared to controls ($p$<0.001, 95%-C.I.=[1.49, 1.91]; $p$<0.001, 95%-C.I.=[0.75, 1.02] respectively). Additionally, EOAD showed a significantly higher tau-PET uptake compared to LOAD ($p$<0.001, 95%-C.I.=[-0.96, -0.54]).

#### Differences in co-pathologies in EO- vs. LOAD

**sTable 5.** AD pathologies, co-pathologies, and cognitive measures of the sample.

| | YCU | OCU | EOAD | LOAD | Total | pFDR YCU-EOAD | pFDR OCU-LOAD | pFDR EOAD-LOAD |
|---|---|---|---|---|---|---|---|---|
| **N** | 188 | 151 | 41 | 154 | 534 | - | - | - |
| **CSF Aβ42/40 ratio** | 1.02±0.13 | 0.99±0.14 | 0.46±0.09 | 0.47±0.11 | 0.81±0.29 | **<.001** | **<.001** | .655 |
| **MTL tau-PET** | 1.02±0.26 | 1.21±0.26 | 2.67±0.59 | 2.67±0.65 | 1.67±0.87 | **<.001** | **<.001** | .975 |
| **Amygdala tau-PET** | 0.82±0.11 | 0.87±0.11 | 2.23±0.67 | 2.08±0.61 | 1.31±0.72 | **<.001** | **<.001** | .207 |
| **EBM-II tau-PET** | 1.04±0.10 | 1.10±0.10 | 2.57±0.92 | 2.18±0.73 | 1.51±0.76 | **<.001** | **<.001** | **.006** |
| **EBM-III tau-PET** | 0.99±0.10 | 1.07±0.11 | 3.01±1.25 | 1.84±0.64 | 1.42±0.77 | **<.001** | **<.001** | **<.001** |
| **EBM-IV tau-PET** | 0.88±0.10 | 0.93±0.10 | 1.68±0.69 | 1.31±0.41 | 1.08±0.40 | **<.001** | **<.001** | **<.001** |
| **EBM-V tau-PET** | 0.94±0.09 | 1.00±0.10 | 1.61±0.61 | 1.24±0.30 | 1.10±0.32 | **<.001** | **<.001** | **<.001** |
| **WMH vol.** | 3320±2200 | 8140±6020 | 5670±3720 | 9690±6950 | 6690±5830 | **<.001** | .085 | **<.001** |
| **WMH vol.** dich. | 13 (6.9) | 69 (45.7) | 11 (26.8) | 80 (51.9) | 173 (32.4) | **<.001** | .277 | **.004** |
| **aHC/PHC ratio +** | 23 (12.2) | 3 (2.0) | 15 (36.6) | 44 (28.6) | 85 (15.9) | **<.001** | **<.001** | .333 |

Continuous variables are displayed as mean±SD. Categorical variables are displayed as n (%). P-values are FDR adjusted. Aβ positivity: <.08 on CSF Aβ42/40 ratio.

Abbreviations: Aβ=amyloid-beta; aHC=anterior hippocampus; CSF=cerebrospinal fluid; dich=dichotomized variable; EBM=event-based modeling; EOAD=early-onset cognitive impairment; FDR=false-discovery rate adjusted p-values; HC=Hippocampus; ROI=region of interest; LOAD=late-onset cognitive impairment; OCU=older cognitively unimpaired controls; PHC=parahippocampal cortex; SD=standard deviation; YCU=younger cognitively unimpaired controls; WMH=white matter hyperintensities.

**sTable 6.** AD pathologies, co-pathologies, and cognitive measures of the sample with comparisons between controls and AD groups adjusted for age.

| | YCU | OCU | EOAD | LOAD | pFDR YCU-EOAD | pFDR OCU-LOAD |
|---|---|---|---|---|---|---|
| **N** | 188 | 151 | 41 | 154 | - | - |
| **CSF Aβ42/40 ratio** | 1.02±0.13 | 0.99±0.14 | 0.46±0.09 | 0.47±0.11 | **<.001** | **<.001** |
| **MTL tau-PET** | 1.02±0.26 | 1.21±0.26 | 2.67±0.59 | 2.67±0.65 | **<.001** | **<.001** |
| **Amygdala tau-PET** | 0.82±0.11 | 0.87±0.11 | 2.23±0.67 | 2.08±0.61 | **<.001** | **<.001** |
| **EBM-II tau-PET** | 1.04±0.10 | 1.10±0.10 | 2.57±0.92 | 2.18±0.73 | **<.001** | **<.001** |
| **EBM-III tau-PET** | 0.99±0.10 | 1.07±0.11 | 3.01±1.25 | 1.84±0.64 | **<.001** | **<.001** |
| **EBM-IV tau-PET** | 0.88±0.10 | 0.93±0.10 | 1.68±0.69 | 1.31±0.41 | **<.001** | **<.001** |
| **EBM-V tau-PET** | 0.94±0.09 | 1.00±0.10 | 1.61±0.61 | 1.24±0.30 | **<.001** | **<.001** |
| **WMH vol.** | 3320±2200 | 8140±6020 | 5670±3720 | 9690±6950 | **<.001** | .033 |
| **WMH vol.** dich. | 13 (6.9) | 69 (45.7) | 11 (26.8) | 80 (51.9) | **<.001** | .262 |
| **aHC/PHC ratio +** | 23 (12.2) | 3 (2.0) | 15 (36.6) | 44 (28.6) | **<.001** | **<.001** |

Continuous variables are displayed as mean±SD. Categorical variables are displayed as n (%). P-values are FDR adjusted. Aβ positivity: <.08 on CSF Aβ42/40 ratio.

Abbreviations: Aβ=amyloid-beta; aHC=anterior hippocampus; CSF=cerebrospinal fluid; dich=dichotomized variable; EOAD=early-onset cognitive impairment, ROI=region of interest; LOAD=late-onset cognitive impairment; OCU=older cognitively unimpaired controls; PHC=parahippocampal cortex; SD=standard deviation; YCU=younger cognitively unimpaired controls.

**sFigure 14.** Group comparison for the dichotomized white matter hyperintensity volumes.

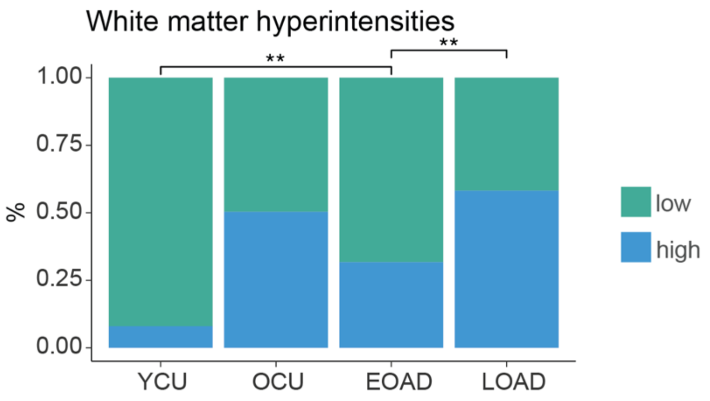

Significant differences are shown for FDR-corrected p-values.
Abbreviations: EOAD=early-onset Alzheimer's Disease; LOAD=late-onset Alzheimer's disease; OCU=older controls; YCU=younger controls.

**sFigure 15.** Group comparison for tau-PET uptake in neocortical composite regions.

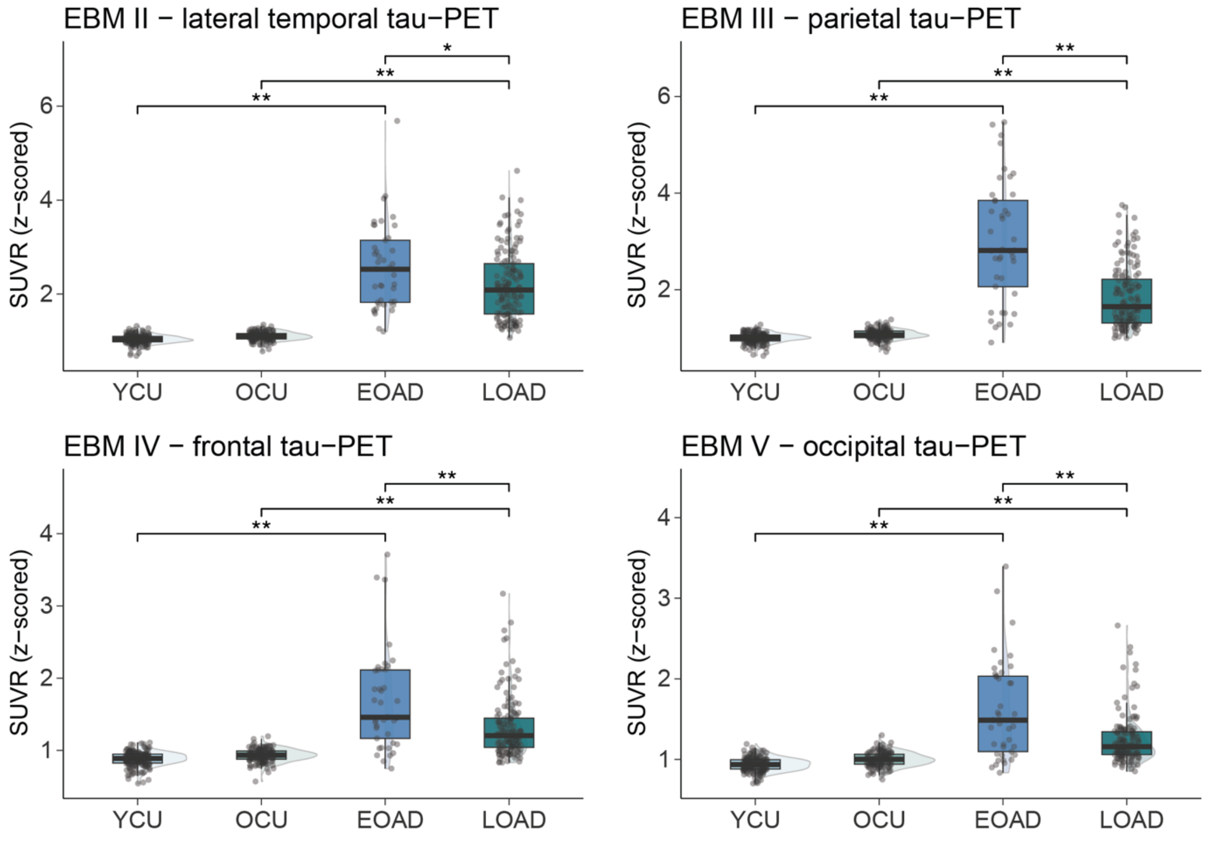

Separate ANOVAs were performed for each comparison. Significant differences are shown for FDR-corrected p-values. A: additional group comparisons for neocortical tau-PET uptake and dichotomized white matter hyperintensity volume. B: exploratory analyses investigating the LEADS signature for thickness and tau-PET uptake. LEADS signature based on Touroutoglou et al. (2023).
Abbreviations: EOAD=early-onset Alzheimer's Disease; LOAD=late-onset Alzheimer's disease; LEADS=Longitudinal Early-Onset Alzheimer's Disease Study; PET=positron emission tomography; SUVR=standardized uptake value ratio; OCU=older controls; YCU=younger controls.

#### Associations between (co-)pathologies and structural measures within amnestic EOAD

**sFigure 16.** Associations between (co-)pathologies of interest with structural measures.

**A**

| | tau-PET SUVR | CSF Aβ42/40 | WMH vol | aHC/pHC ratio |
|---|---|---|---|---|
| Total Hippocampus | 0.06 | 0.08 | 0.16 | –0.63 |
| Entorhinal cortex | –0.15 | –0.14 | –0.01 | –0.33 |
| Parahippocampal cortex | 0.06 | –0.19 | –0.24 | 0.16 |

p<.05

**B**

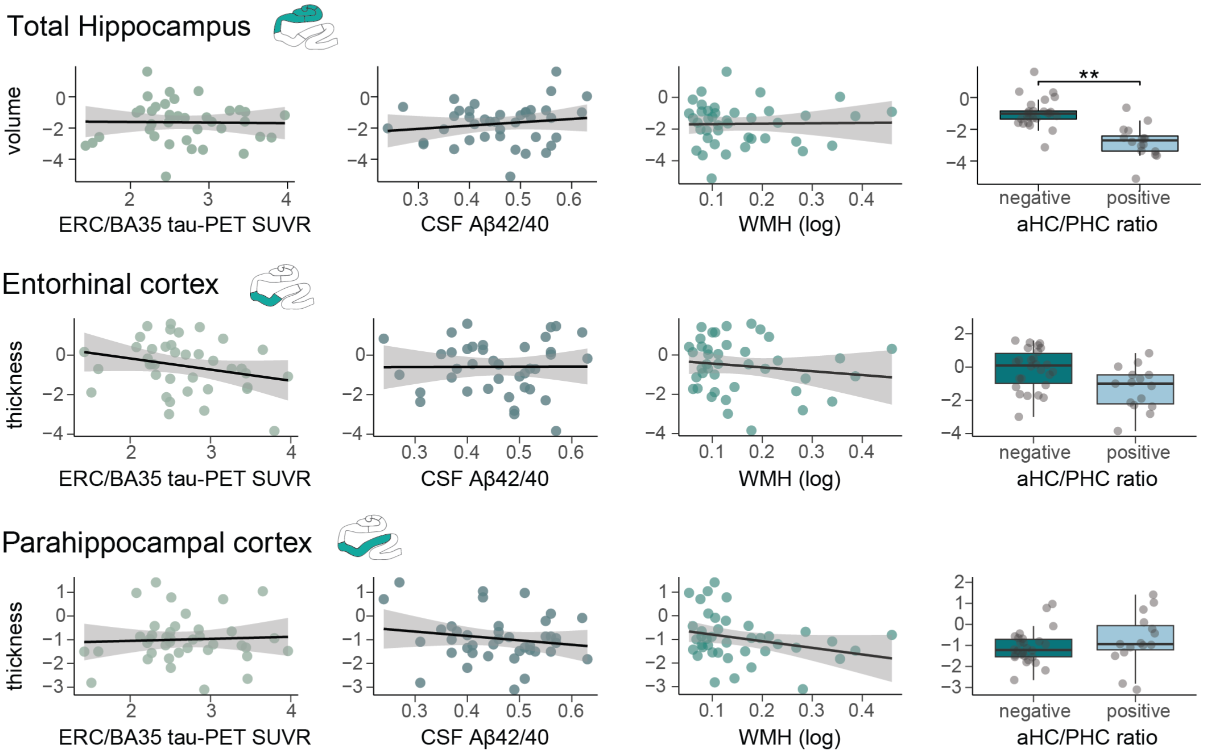

Exploratory linear regressions were performed focusing on EOAD only. A: shows the standardized beta coefficients of the linear regression models. Significant associations are shown with FDR-corrected p-values (colored cells are $p_{FDR}$<.05). B: shows scatterplots and boxplots of the investigated associations.

Abbreviations: BA35=Brodmann area 35; CSF=cerebrospinal fluid; EOAD=early-onset Alzheimer's Disease; ERC=entorhinal cortex; LOAD=late-onset Alzheimer's disease; NEG=negative; PET=positron emission tomography; POS=positive; SUVR=standardized uptake value ratio; OCU=older controls; YCU=younger controls.

### Cognitive performance in amnestic EOAD

**sTable 7.** Comparisons of cognitive performance across the groups.

| | YCU | OCU | EOAD | LOAD | Total | p-value YCU-EOAD | p-value OCU-LOAD | p-value EOAD-LOAD |
|---|---|---|---|---|---|---|---|---|
| **N** | 188 | 151 | 41 | 154 | 534 | - | - | - |
| **ADAS delayed** | 1.89±1.49 | 2.98±1.82 | 8.02±1.44 | 8.53±1.45 | 4.59±3.34 | **<.001** | **<.001** | .291 |
| **Animal fluency** | 26.3±5.74 | 23.0±4.99 | 17.2±5.37 | 13.9±4.85 | 21.1±7.36 | **<.001** | **<.001** | **.004** |
| **BNT-15** | 14.3±0.99 | 13.7±1.42 | 12.7±2.56 | 10.9±2.83 | 13.0±2.40 | **<.001** | **<.001** | **<.001** |
| **VOSP cube** | 9.73±0.67 | 9.56±0.98 | 7.84±2.85 | 8.41±2.14 | 9.18±1.65 | **<.001** | **<.001** | .197 |
| **SDM** | 48.3±9.31 | 36.7±7.93 | 26.5±12.9 | 24.6±8.14 | 36.9±13.2 | **<.001** | **<.001** | .662 |
| **TMT-B** | 70.4±24.4 | 100±45.3 | 227±139 | 250±124 | 138±111 | **<.001** | **<.001** | .466 |

Continuous variables are displayed as mean±SD. Categorical variables are displayed as n (%). [a] individuals who reported an age-of-onset under 65 were included in the EOAD group. [b] Aβ positivity: <.08 on CSF Aβ42/40 ratio.
Abbreviations: Aβ=amyloid-beta; CSF=cerebrospinal fluid; EOAD=early-onset cognitive impairment, LOAD=late-onset cognitive impairment; OCU=older cognitively unimpaired controls; SD=standard deviation; YCU=younger cognitively unimpaired controls.

**sFigure 17.** Associations between all cognitive test scores with structural MRI measures.

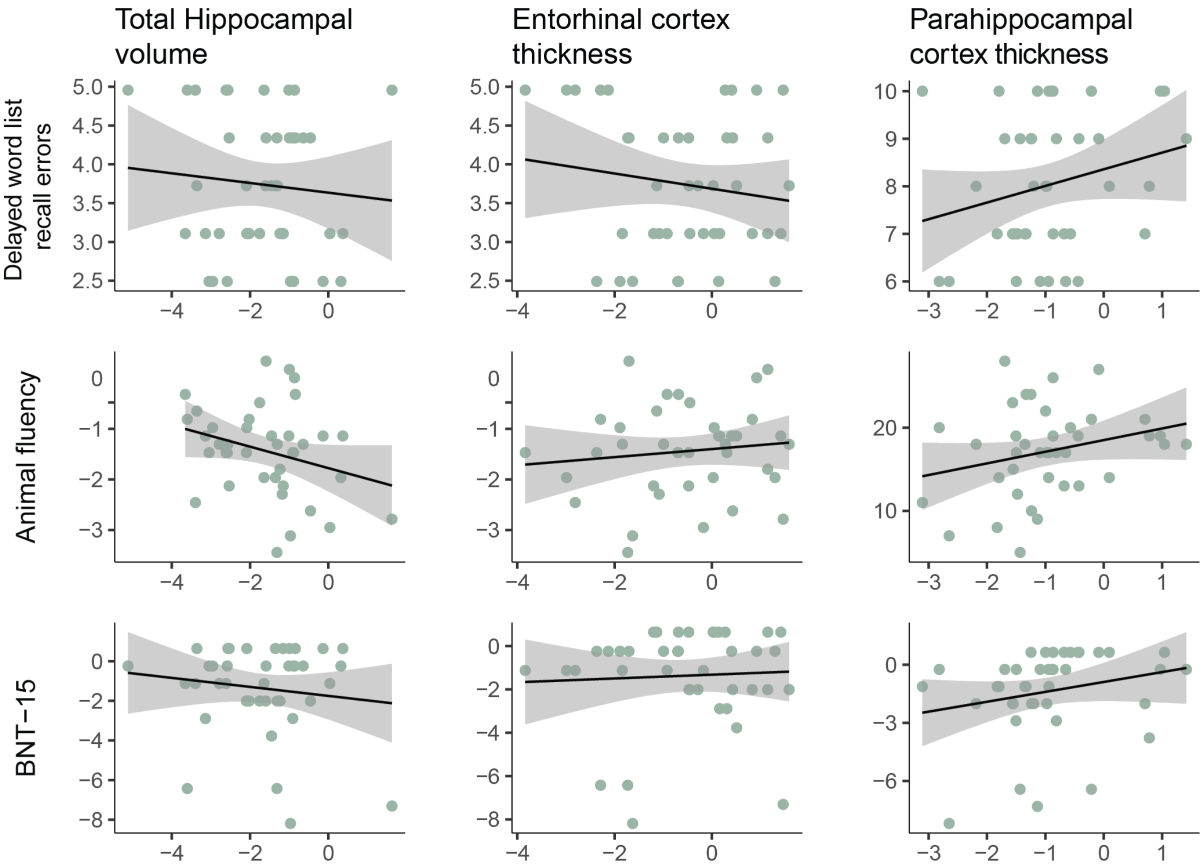

Linear regressions were performed using only the EOAD group (n=41) and including age, sex, and education as covariates. None of the associations were statistically significant.

### Comparison between amnestic and non-amnestic EO- and LOAD

#### Demographics

**sTable 8.** Characteristics of the sample including non-amnestic Alzheimer's disease cases.

| | YCU | OCU | EOAD | naEOAD | LOAD | naLOAD | Total |
|---|---|---|---|---|---|---|---|
| **N** | 188 | 151 | 41 | 7 | 154 | 16 | 557 |
| **Diagnosis** (CU/MCI/AD) | 188/0/0 | 151/0/0 | 0/16/25 | 0/3/4 | 0/65/89 | 0/10/6 | 339/95/123 |
| **Sex** (female) | 103 (54.8) | 99 (65.5) | 20 (48.8) | 3 (42.9) | 82 (53.2) | 7 (43.8) | 314 (56.4) |
| **Age** | 58.6±4.89 | 77.3±3.38 | 61.0±4.82 | 61.6±3.98 | 76.2±3.92 | 77.4±3.98 | 69.3±9.72 |
| Range | 51.0 – 69.0 | 70.3 – 85.0 | 50.9 – 69.4[a] | 56.0 – 66.1 | 70.1 – 85.1 | 71.3 – 85.4 | 50.9 – 85.4 |
| **Education** (years) | 13.2±3.12 | 12.4±3.74 | 14.1±3.33 | 13.6±2.82 | 12.5±4.79 | 12.8±4.09 | 12.8±3.85 |
| Missing | 2 (1.1) | 0 (0.0) | 1 (2.4) | 0 (0.0) | 6 (3.9) | 0 (0.0) | 9 (1.6) |
| ***APOE-ε4* allele**[b] | 85 (45.2) | 29 (19.2) | 25 (61.0) | 5 (71.4) | 114 (74.0) | 9 (56.3) | 267 (47.9) |
| **CSF Aβ42/40 +** | 0 (0.0) | 0 (0.0) | 41 (100) | 7 (100) | 154 (100) | 16 (100) | 218 (39.1) |

Continuous variables are displayed as mean±SD. Categorical variables are displayed as n (%). [a] Aβ positivity: <.08 on CSF Aβ42/40 ratio. Abbreviations: Aβ=amyloid-beta; aHC=anterior hippocampus; CSF=cerebrospinal fluid; EOAD=amnestic early-onset cognitive impairment; ROI=region of interest; LOAD=amnestic late-onset cognitive impairment; naEOAD=non-amnestic early-onset AD; naLOAD=non-amnestic late-onset AD; OCU=older cognitively unimpaired controls; PHC=parahippocampal cortex; SD=standard deviation; YCU=younger cognitively unimpaired controls.

**sTable 9.** Comparison between groups on structural MRI measures including non-amnestic AD groups.

| | YCU vs. EOAD | | YCU vs. naEOAD | | OCU vs. LOAD | | OCU vs. naLOAD | | EOAD vs. LOAD | | naEOAD vs. naLOAD | | EOAD vs. naEOAD | | LOAD vs. naLOAD | |
|---|---|---|---|---|---|---|---|---|---|---|---|---|---|---|---|---|
| | diff | $p_{FDR}$ | diff | $p_{FDR}$ | diff | $p_{FDR}$ | diff | $p_{FDR}$ | diff | $p_{FDR}$ | diff | $p_{FDR}$ | diff | $p_{FDR}$ | diff | $p_{FDR}$ |
| SUB | 1.617 | **<.001** | 0.454 | 0.215 | 1.249 | **<.001** | 1.136 | **<.001** | 0.502 | **0.008** | 1.551 | **<.001** | -1.162 | **0.011** | -0.113 | 0.722 |
| DG | 1.043 | **<.001** | 0.378 | 0.419 | 0.993 | **<.001** | 0.674 | **0.022** | 0.380 | 0.066 | 0.726 | 0.154 | -0.665 | 0.201 | -0.319 | 0.305 |
| CA1 | 1.124 | **<.001** | 0.049 | 0.877 | 1.122 | **<.001** | 0.509 | 0.065 | 0.387 | 0.066 | 0.849 | 0.107 | -1.075 | **0.041** | -0.613 | **0.041** |
| ERC | 0.808 | **<.001** | -0.235 | 0.804 | 1.598 | **<.001** | 0.438 | **0.093** | 0.779 | **0.006** | 0.662 | 0.397 | -1.043 | 0.109 | -1.160 | **0.007** |
| BA35 | 1.698 | **<.001** | 0.644 | 0.121 | 1.196 | **<.001** | 0.827 | **0.003** | 0.409 | 0.088 | 1.095 | 0.072 | -1.054 | **0.022** | -0.368 | 0.355 |
| BA36 | 0.608 | **<.001** | 0.272 | 0.411 | 0.601 | **<.001** | 0.464 | **0.014** | 0.337 | 0.092 | 0.536 | 0.263 | -0.336 | 0.543 | -0.138 | 0.793 |
| PHC | 1.035 | **<.001** | 0.887 | **0.039** | 0.913 | **<.001** | 0.991 | **<.001** | 0.677 | **0.001** | 0.902 | 0.112 | -0.148 | 0.793 | 0.077 | 0.882 |
| Total HC | 1.684 | **<.001** | 0.369 | 0.373 | 1.547 | **<.001** | 0.775 | **0.002** | 0.499 | **0.020** | 1.043 | 0.055 | -1.316 | **0.037** | -0.772 | **0.010** |
| AMY | 1.899 | **<.001** | 0.404 | 0.341 | 1.548 | **<.001** | 0.892 | **0.002** | 0.366 | 0.207 | 1.205 | 0.072 | -1.495 | **0.048** | -0.656 | 0.094 |
| LT | 1.314 | **<.001** | 1.061 | **0.001** | 1.036 | **<.001** | 1.013 | **<.001** | 0.381 | **0.048** | 0.612 | 0.280 | -0.254 | 0.603 | -0.023 | 0.902 |
| LP | 1.536 | **<.001** | 1.470 | **0.000** | 0.754 | **<.001** | 0.825 | **0.006** | -0.114 | 0.676 | 0.022 | 0.947 | -0.066 | 0.888 | 0.070 | 0.888 |
| MP | 1.439 | **<.001** | 1.249 | **0.003** | 0.849 | **<.001** | 0.776 | **0.014** | 0.049 | 0.804 | 0.166 | 0.798 | -0.190 | 0.782 | -0.073 | 0.793 |
| FL | 0.510 | **0.002** | 0.454 | 0.263 | 0.689 | **<.001** | 0.826 | **0.008** | 0.461 | **0.023** | 0.654 | 0.227 | -0.056 | 0.877 | 0.137 | 0.804 |
| OL | 0.371 | 0.054 | 0.047 | 0.892 | 0.276 | **0.044** | 0.190 | 0.551 | 0.157 | 0.521 | 0.395 | 0.504 | -0.324 | 0.646 | -0.086 | 0.804 |

Positive mean differences indicate higher values in the group listed first; negative mean differences indicate lower values in the group listed first. All analyses were adjusted for sex. All p-values are FDR adjusted.
Abbreviations: aHC=anterior hippocampus; AMY=amygdala; BA=Brodmann area; CA1=cornu ammonis 1; DG=dentate gyrus; diff=mean difference; ERC=entorhinal cortex; EOAD=amnestic early-onset cognitive impairment; FL=frontal cortex; HC=hippocampus; LOAD=amnestic late-onset cognitive impairment; LT=lateral temporal; LP=lateral parietal; MP=medial parietal; naEOAD=non-amnestic early-onset AD; naLOAD=non-amnestic late-onset AD; OCU=older cognitively unimpaired controls; OL=occipital cortex; PHC=parahippocampal cortex; SUB=subiculum; YCU=younger cognitively unimpaired controls.

**sTable 10.** Comparison between groups on AD biomarkers and co-pathologies and cognitive measures.

| | **YCU-EOAD** | | **YCU-naEOAD** | | **OCU-LOAD** | | **OCU-naLOAD** | | **EOAD-LOAD** | | **naEOAD-naLOAD** | | **EOAD-naEOAD** | | **LOAD-naLOAD** | |
|---|---|---|---|---|---|---|---|---|---|---|---|---|---|---|---|---|
| | diff | $p_{FDR}$ | diff | $p_{FDR}$ | diff | $p_{FDR}$ | diff | $p_{FDR}$ | diff | $p_{FDR}$ | diff | $p_{FDR}$ | diff | $p_{FDR}$ | diff | $p_{FDR}$ |
| CSF Aβ42/40 | 0.561 | **<.001** | 0.533 | **<.001** | 0.527 | **<.001** | 0.575 | **<.001** | -0.009 | .727 | 0.067 | .170 | -0.028 | .638 | 0.048 | .161 |
| ERC/BA35 tau-PET SUVR | -1.646 | **<.001** | -1.177 | **<.001** | -1.455 | **<.001** | -1.168 | **<.001** | 0.001 | .982 | -0.182 | .623 | 0.469 | .148 | 0.286 | .182 |
| EBM II tau-PET SUVR | -1.538 | **<.001** | -1.255 | **<.001** | -1.081 | **<.001** | -1.045 | **<.001** | 0.390 | **.008** | 0.144 | .831 | 0.283 | .601 | 0.036 | .919 |
| EBM III tau-PET SUVR | -2.011 | **<.001** | -1.399 | **<.001** | -0.772 | **<.001** | -0.740 | **<.001** | 1.167 | **<.001** | 0.586 | .346 | 0.612 | .365 | 0.032 | .923 |
| EBM IV tau-PET SUVR | -0.799 | **<.001** | -0.634 | **<.001** | -0.382 | **<.001** | -0.484 | **<.001** | 0.366 | **<.001** | 0.099 | .892 | 0.165 | .669 | -0.102 | .558 |
| EBM V tau-PET SUVR | -0.676 | **<.001** | -0.489 | **<.001** | -0.242 | **<.001** | -0.243 | **<.001** | 0.371 | **<.001** | 0.183 | .553 | 0.187 | .601 | -0.001 | .919 |
| Amygdala tau-PET SUVR | -1.404 | **<.001** | -0.673 | **<.001** | -1.206 | **<.001** | -0.624 | **<.001** | 0.148 | .270 | -0.001 | .998 | 0.731 | **.032** | 0.582 | **.002** |
| WMH volume | -0.065 | **<.001** | -0.052 | .063 | -0.041 | .120 | -0.163 | **.001** | -0.111 | **<.001** | -0.247 | .061 | 0.013 | .828 | -0.123 | **.034** |
| Delayed word list recall | -3.780 | **<.001** | -1.563 | **<.001** | -3.423 | **<.001** | -1.014 | **.003** | -0.313 | .176 | -0.121 | .638 | 2.217 | **<.001** | 2.409 | **<.001** |
| Animal fluency | 1.497 | **<.001** | 1.362 | **.001** | 1.498 | **<.001** | 1.610 | **<.001** | 0.544 | **.001** | 0.791 | .182 | -0.135 | .842 | 0.112 | .663 |
| BNT-15 | 1.411 | **<.001** | 1.925 | **<.001** | 2.481 | **<.001** | 1.837 | **<.001** | 1.616 | **.004** | 0.457 | .842 | 0.514 | .727 | -0.644 | .545 |
| VOSP Cube | 2.935 | **<.001** | 1.441 | **.010** | 1.783 | **<.001** | 2.220 | **<.001** | -0.896 | .148 | 1.035 | .736 | -1.493 | .566 | 0.437 | .638 |
| Symbol Digit Modalities | 1.815 | **<.001** | 2.111 | **<.001** | 1.017 | **<.001** | 1.165 | **<.001** | 0.164 | .553 | 0.017 | .924 | 0.296 | .540 | 0.148 | .566 |
| TMT-B | -4.385 | **<.001** | -4.752 | **<.001** | -4.189 | **<.001** | -4.378 | **<.001** | -0.643 | .727 | -0.465 | .831 | -0.367 | .878 | -0.189 | .895 |
| | OR | $p_{FDR}$ | OR | $p_{FDR}$ | OR | $p_{FDR}$ | OR | $p_{FDR}$ | OR | $p_{FDR}$ | OR | $p_{FDR}$ | OR | $p_{FDR}$ | OR | $p_{FDR}$ |
| aHC/PHC ratio + | 4.575 | **<.001** | 3.121 | .288 | 20.073 | **<.001** | 2.658 | .553 | 0.685 | .437 | 0.143 | .274 | 0.716 | .801 | 5.637 | .161 |

All analyses were adjusted for sex. All p-values are FDR adjusted.

Abbreviations: BA35=Brodmann area 35; BNT-15=Boston Naming Test-15; diff=mean difference; ERC=entorhinal cortex; EOAD=amnestic early-onset cognitive impairment; LOAD=amnestic late-onset cognitive impairment; naEOAD=non-amnestic early-onset AD; naLOAD=non-amnestic late-onset AD; OCU=older cognitively unimpaired controls; OR=odds ratui; YCU=younger cognitively unimpaired controls.

**sFigure 18.** Boxplots showing significant differences of comparisons with the non-amnestic and amnestic AD groups for medial temporal lobe structural MRI measures.

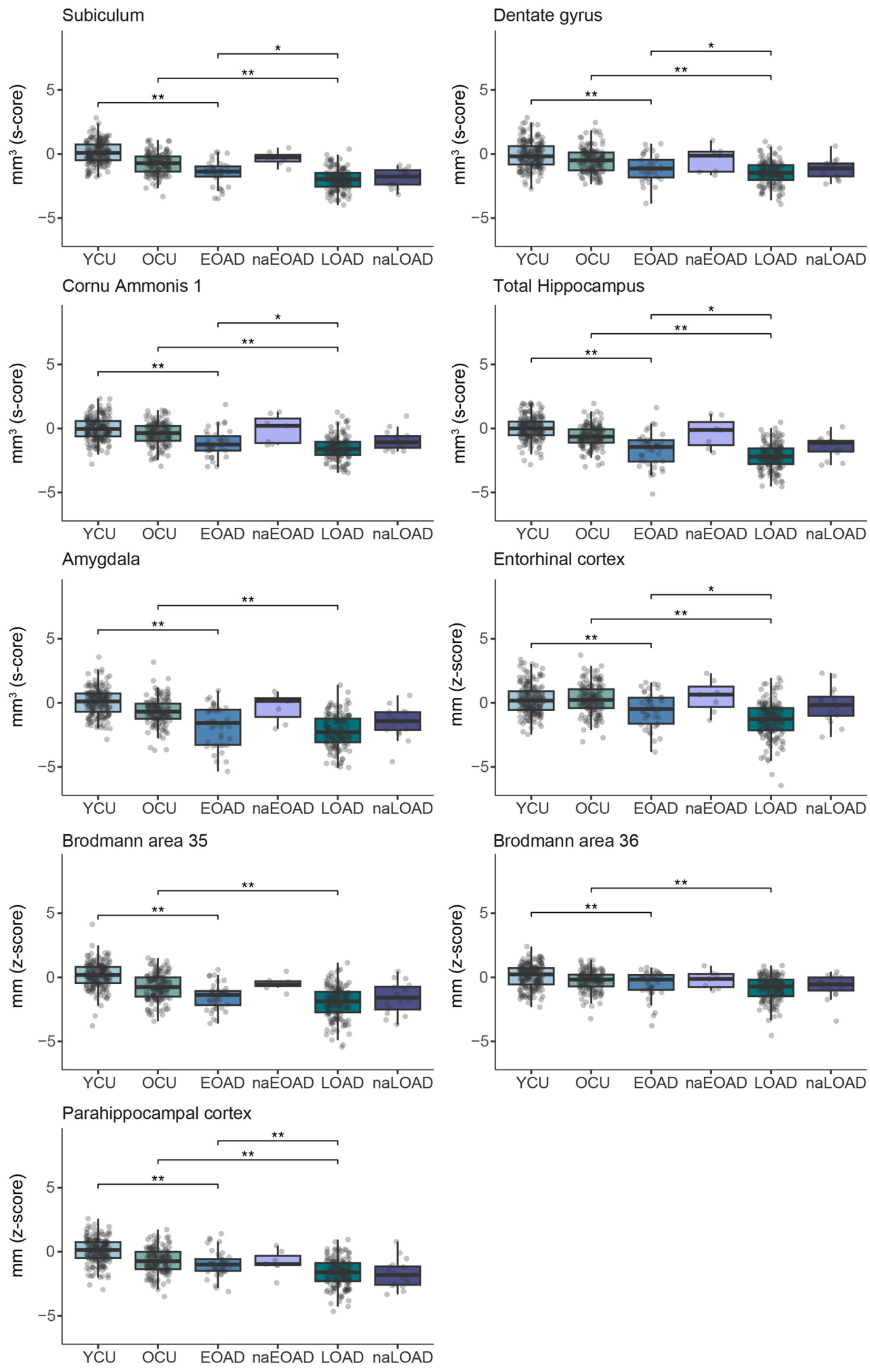

ANOVAs were performed for each comparison. Significant differences are shown for FDR-corrected p-values. Abbreviations: EOAD=amnestic early-onset Alzheimer's Disease; LOAD=amnestic late-onset Alzheimer's disease; naEOAD=non-amnestic early-onset AD; naLOAD=non-amnestic late-onset AD; OCU=older controls; YCU=younger controls.

**sFigure 19.** Boxplots showing significant differences of comparisons with the non-amnestic and amnestic AD groups for the neocortical structural measures.

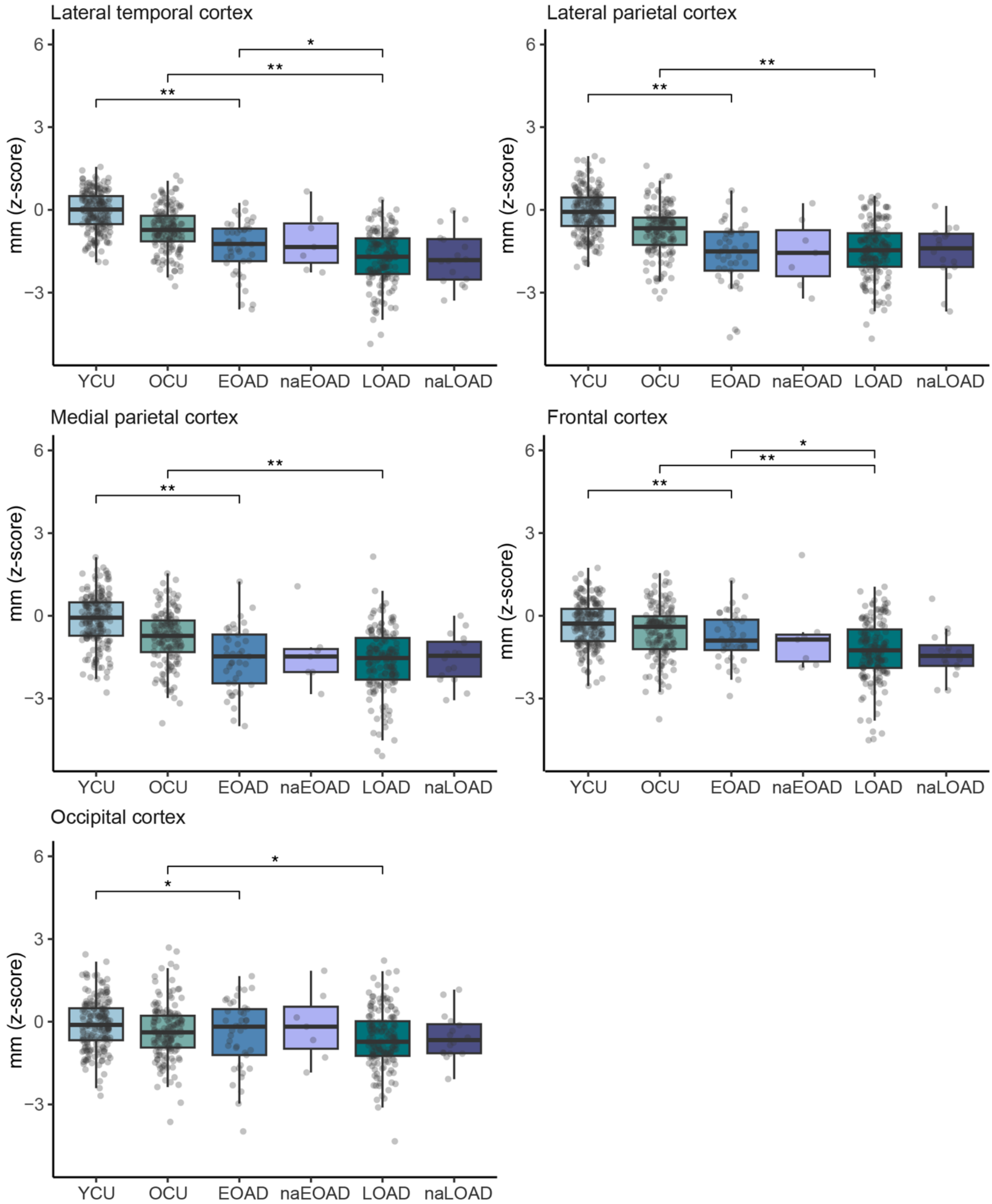

ANOVAs were performed for each comparison. Significant differences are shown for FDR-corrected p-values. Abbreviations: EOAD=amnestic early-onset Alzheimer's Disease; LOAD=amnestic late-onset Alzheimer's disease; naEOAD=non-amnestic early-onset AD; naLOAD=non-amnestic late-onset AD; OCU=older controls; YCU=younger controls.

**sFigure 20.** Boxplots showing significant differences of comparisons with the non-amnestic and amnestic AD groups for the significant pathologies and cognitive measures.

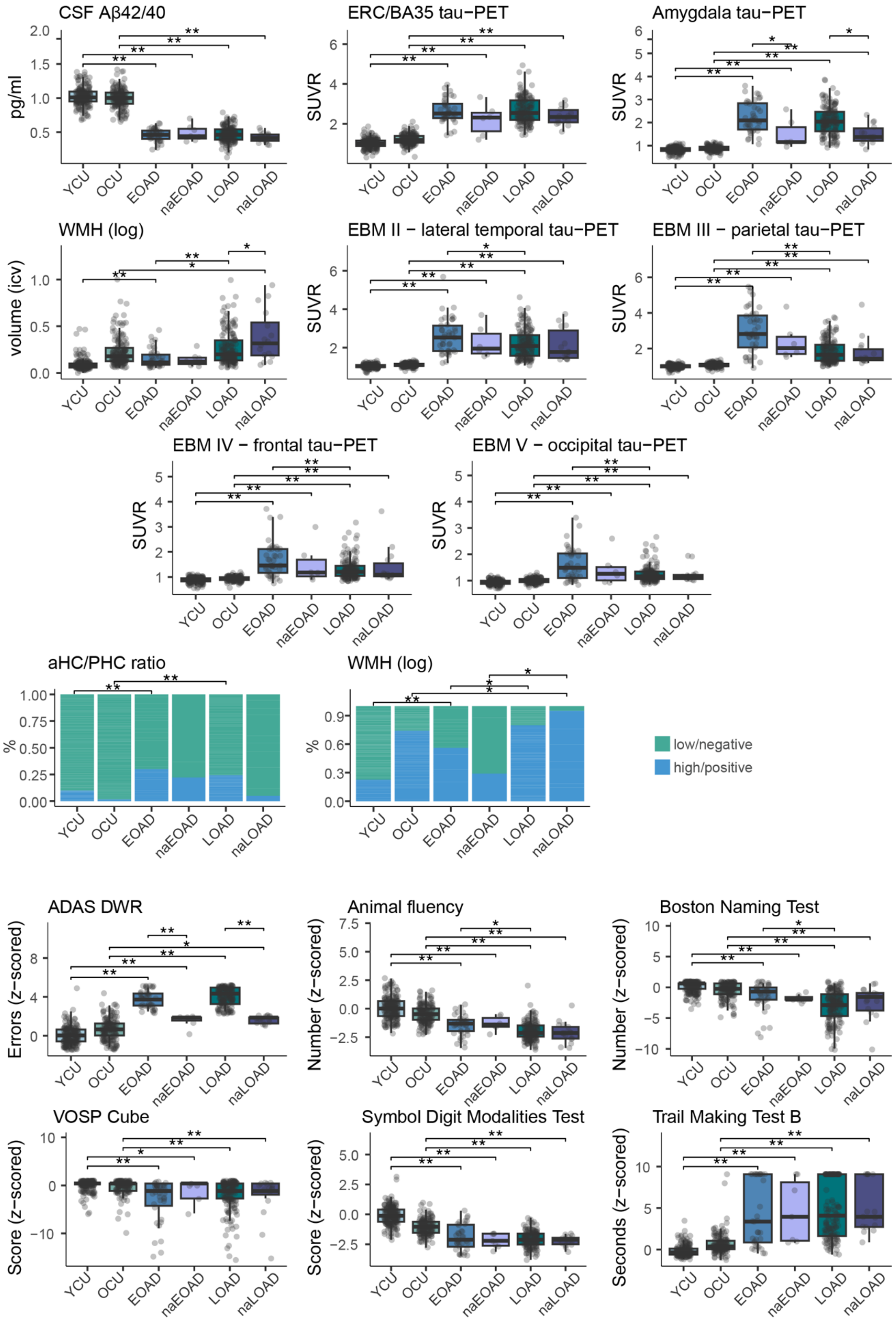

ANOVAs were performed for each comparison. Significant differences are shown for FDR-corrected p-values. Abbreviations: EOAD=amnestic early-onset Alzheimer's Disease; LOAD=amnestic late-onset Alzheimer's disease; naEOAD=non-amnestic early-onset AD; naLOAD=non-amnestic late-onset AD; OCU=older controls; YCU=younger controls.
